# Disentangling semantic composition and semantic association in the left temporal lobe

**DOI:** 10.1101/2020.08.17.254482

**Authors:** Jixing Li, Liina Pylkkänen

**Author notes:** corresponding author Jixing Li. Contributions: designed research, performed research, analyzed data, wrote the paper. Contributions: designed research, wrote the paper.

## Abstract

Although composing two words into a complex representation (e.g., “coffee cake”) is conceptually different from forming associations between a pair of words (e.g., “coffee, cake”), the brain regions supporting semantic composition have also been implicated for associative encoding. Here, we adopted a two-word magnetoencephalography (MEG) paradigm which varies compositionality (“French/Korean cheese” vs. “France/Korea cheese”) and strength of association (“France/French cheese” vs. “Korea/Korean cheese”) between the two words. We collected MEG data while 42 English speakers (24 females) viewed the two words successively in the scanner, and we applied both univariate regression analyses and multivariate pattern classification to the source estimates of the two words. We show that the left anterior and middle temporal lobe (LATL; LMTL) are distinctively modulated by semantic composition and semantic association. Specifically, the LATL is mostly sensitive to high-association compositional phrases, while the LMTL responds more to low-association compositional phrases. Pattern-based directed connectivity analyses further revealed a continuous information flow from the anterior to the middle temporal region, suggesting that the integration of adjective and noun properties originated earlier in the LATL is consistently delivered to the LMTL when the complex meaning is newly encountered. Taken together, our findings shed light into a functional dissociation within the left temporal lobe for compositional and distributional semantic processing.

**Significance Statement:** Prior studies on semantic composition and associative encoding have been conducted independently within the subfields of language and memory, and they typically adopt similar two-word experimental paradigms. However, no direct comparison has been made on the neural substrates of the two processes. The current study relates the two streams of literature, and appeals to audiences in both subfields within cognitive neuroscience. Disentangling the neural computations for semantic composition and association also offers insight into modeling compositional and distributional semantics, which has been the subject of much discussion in Natural Language Processing and cognitive science.

## Introduction

When we hear a pair of words such as “coffee” and “cake”, we could form a complex meaning of a coffee-flavored cake, or recall an experience where coffee and cake occurred together. These two processes, though conceptually different, have been localized to similar brain regions in the left temporal lobe. For example, a minimal adjective-noun phrase such as “red boat” elicits increased activity in the left anterior temporal lobe (LATL) compared to non-compositional word lists such as “cup, boat” (Bemis and Pylkkänen, 2011, 2013), and a similar effect has been observed for a language with the reverse word order (Westerlund et al., 2015) and for American Sign Language (Blanco-Elorrieta et al., 2018), suggesting a role of the LATL in conceptual combination. However, the LATL is also activated when forming arbitrary associations between pairs of words such as “ring” and “cheese” (Jang et al., 2017), and shows greater oscillatory responses for word pairs with stronger associations (Teige et al., 2018, 2019). Additionally, a temporary virtual lesion induced by repetitive transcranial magnetic stimulation (rTMS) over the LATL significantly slows synonym judgment times (Pobric et al., 2007), mirroring a core feature of semantic dementia (SD) patients with a focal atrophy of the ATL (Galton et al., 2001; Hodges et al., 1992; Nestor et al., 2006). The LATL is also engaged in human face-name (Wang et al., 2017) and face-location (Nieuwenhuis et al., 2011) associations, and single neuron recordings in the ATL of macaque monkey’s brain suggested that a group of pair-encoding neurons responded only to associated pairs of abstract shapes during the training phase (e.g., Hirabayashi et al., 2014; Sakai and Miyashita, 1991).

Prior studies on semantic composition and associative encoding have been conducted independently within the subfields of language and memory, and they typically adopt similar two-word experimental paradigms. However, no direct comparison has been made on the neural substrates of the two processes. They both involve connecting two elements, albeit in intuitively different ways. The converging neural evidence in the left anterior temporal cortex raises the question of whether they may involve some shared processing routines. This study aims to characterize how effects of composition and association compare to each other and possibly interact in the left temporal lobe. We adopted the same two-word paradigm but crossed compositionality and strength of association between the two words using the adjectival and noun forms of a country word and our world knowledge of food. “France cheese” and “French cheese”, for example, are highly-associated country-food pairs compared to “Korea cheese” and “Korean cheese”. But both “French cheese” and “Korean cheese” are combinatory adjective-noun phrases whereas “France cheese” and “Korea cheese” are merely word lists. This 2 x 2 design (Figure 1) enabled us to separate the effects of association and composition during two-word comprehension, and to examine whether the LATL indeed supports semantic composition or simply tracks association.

**Figure 1.**
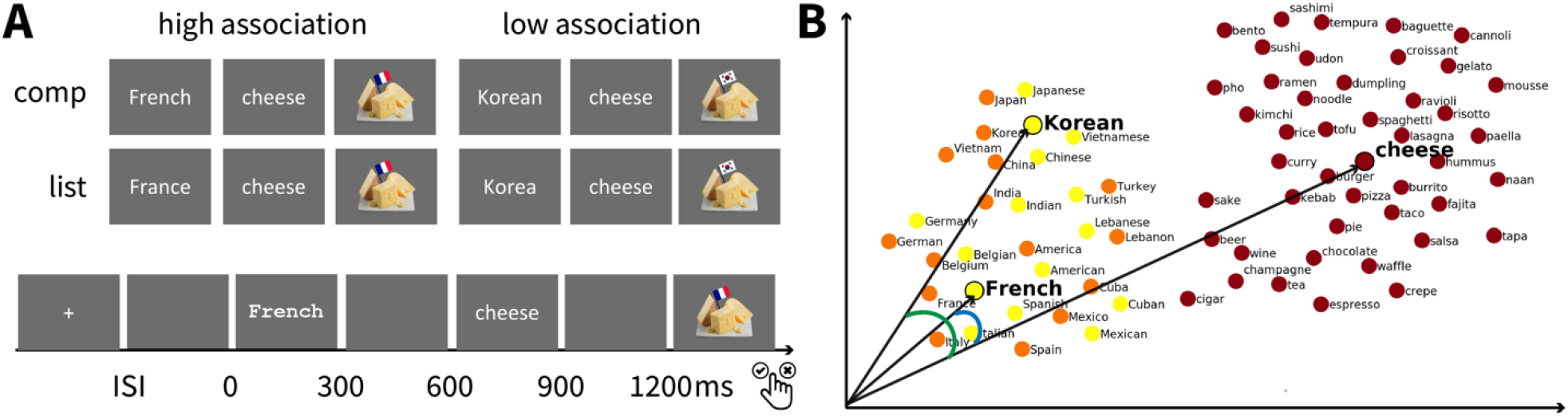
Experimental design. ***A***, Experimental design and trial structure. Our design crossed strength of association (Low vs High) and compositionality (List vs Comp). In each trial, participants indicated whether the target picture matched the preceding words. Half of the target pictures matched and half did not. Activities recorded from 100 ms pre-stimuus onset to 1200 ms post-stimulus onset were analyzed. ***B***, Schematic diagram showing how cosine value between high-dimensional vectors represents semantic similarity. The smaller the angle is between two vectors, the higher the cosine value and semantic similarity. The angle between “French” and “cheese” is smaller than the one between “Korean” and “cheese” because they share more contextual features. The high-dimensional word embeddings were visualized on the 2D scatter using *t*-Distributed Stochastic Neighbor Embedding (t-SNE).

We collected 42 native English speakers’ MEG data while they viewed the two words successively on a screen. We coded association and composition as binary variables and applied a mass univariate regression analysis to source-localized MEG data within a language network.

We then performed a searchlight-based multivariate pattern classification analysis within the same language network. To further understand the information flow between the active brain regions, we conducted a directed connectivity analysis using the representational dissimilarity matrices (RDMs) of the MEG data within the functional regions of interest (fROIs) derived from the regression and the classification analyses.

With this combination of methods, we show that the LATL is more sensitive to the contrast between high and low associative compositional phrases, while the LMTG is mainly driven by the distinction between low-association compositional phrases and low-association lists. Directed connectivity analyses suggest a continuous information flow from the LATL to the LMTG. Taken together, our results provide novel evidence that the LATL and the LMTL are distinctly modulated by semantic composition and semantic association.

## Materials and Methods

### Experimental design

We conducted two variates of experiment, both employed a 2 x 2 design with associative strength (Low, High) and compositionality (Comp, List) as factors. In the first experiment, there are 60 trials for each of the 4 conditions and 60 single word controls consisted of 5 “x”s and the food noun (e.g., “xxxxx cheese”), forming a total of 300 unique trials. In the second experiment, each condition contains 45 trials and 45 single word controls, forming a total of 360 unique trials. The single word trials consisted of length-matched consonant strings and the food noun (e.g., “xjtgbv cheese”).

### Participants

Participants were healthy young adults with normal hearing and normal or corrected-to-normal vison. All strictly qualified as right-handed on the Edinburgh handedness inventory (Oldfield, 1971). They self-identified as native English speakers and gave their written informed consent prior to participation, in accordance with New York University and New York University Abu Dhabi IRB guidelines. 30 volunteers participated in Experiment 1, 9 of them were removed from data analysis: 8 due to excessive movement or drowsiness during MEG recording, and 1 due to bad performance on the behavioral task (less than 75% accuracy). 26 volunteers participated in Experiment 2, 5 of them were removed from further data analysis: 4 due to excessive movement or drowsiness during MEG recording, and 1 due to bad performance on the behavioral task. Therefore, a total of 21 participants (10 females, mean age=21.6 years, SD=4.5) in Experiment 1 and 21 participants (14 females, mean age=23 years, SD=7.7) in Experiment 2 were included in the analyses. No participant attended both experiments. Our sample size was not determined in advance, but it is significantly larger than the typical sample size of around 20–25 participants in prior MEG studies on semantic composition (e.g., Bemis and Pylkkänen, 2011; Westerlund and Pylkkänen, 2014).

Experiment 1 was conducted at the Neuroscience of Language Lab at New York University Abu Dhabi, and Experiment 2 was conducted across research facilities of the Neuroscience of Language Lab at New York University and New York University Abu Dhabi. 6 participants’ data were collected at the New York facility.

### Stimuli

The stimuli comprised of a country adjective/noun and a food noun, with strength of association (Low, High) and compositionality (Comp, List) as factors (Figure 1A). We quantified the strength of association between the country and food word using the cosine similarity score between the GloVe embeddings^17^ of the two words: High associative phrases (e.g., “French/France cheese”) have a cosine similarity greater than 0.3 and low associative pairs (e.g., “Korean/Korea cheese”) have a cosine similarity lower than 0.15 (see Table 1 in Extended Data). Note that, although the noun-noun lists (e.g., “France cheese”) were not syntactically composable, participants may still attempt conceptual composition during the task. To ensure that the list conditions were indeed non-compositional as intended, we included a single-word control condition. The single-word condition consisted of either a “xxxxx” or a length-matched consonant string (e.g., “xjtgbv”) and the critical noun word (i.e., “cheese”). Half of the participants (n=21) saw the “xxxxx” while the other half saw the consonant strings. One ancillary aim of our study was to examine the impact of these two different types of visual baselines, but they patterned the same as regards our main manipulation (see Figure 7)

**Table 1.**
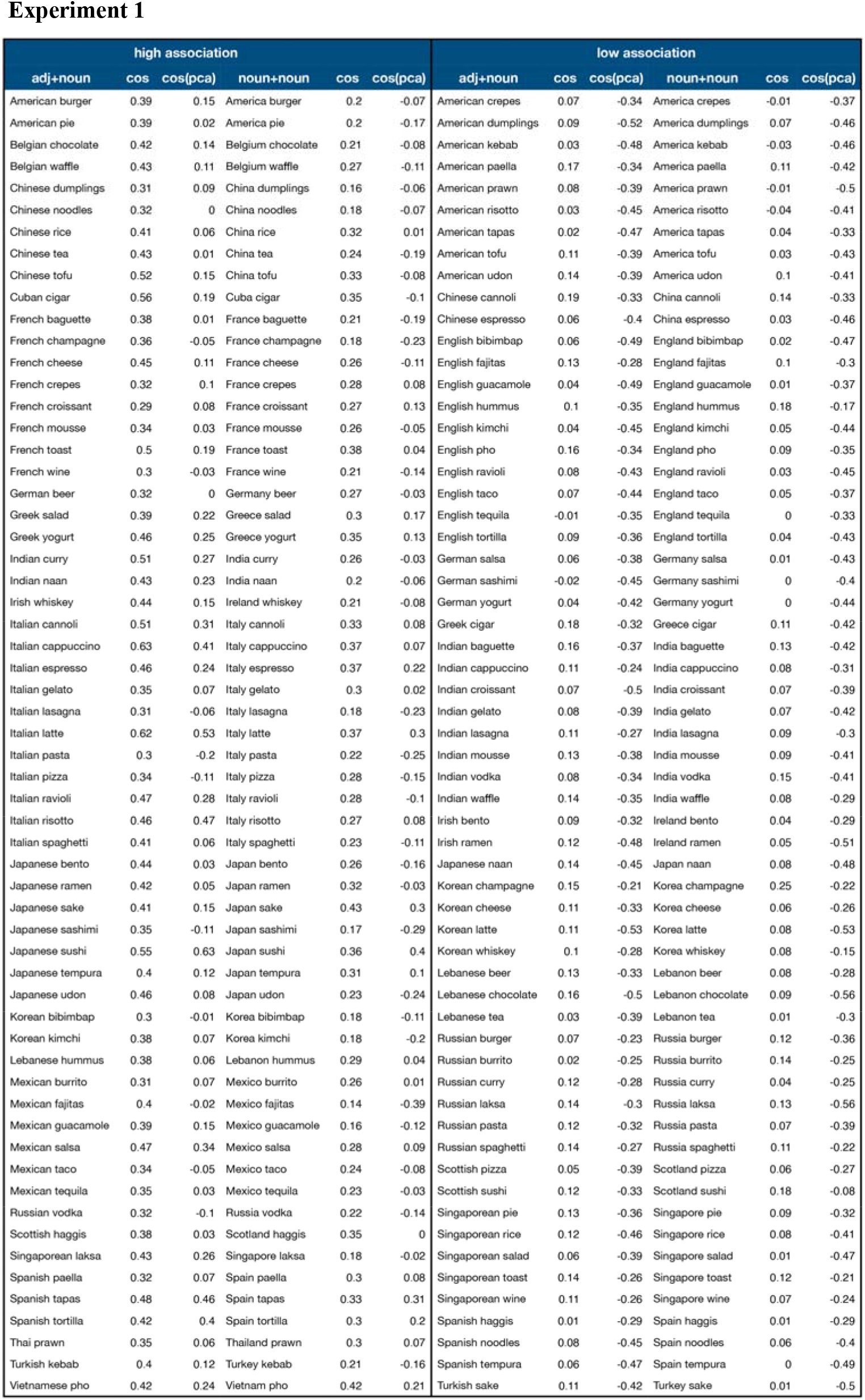
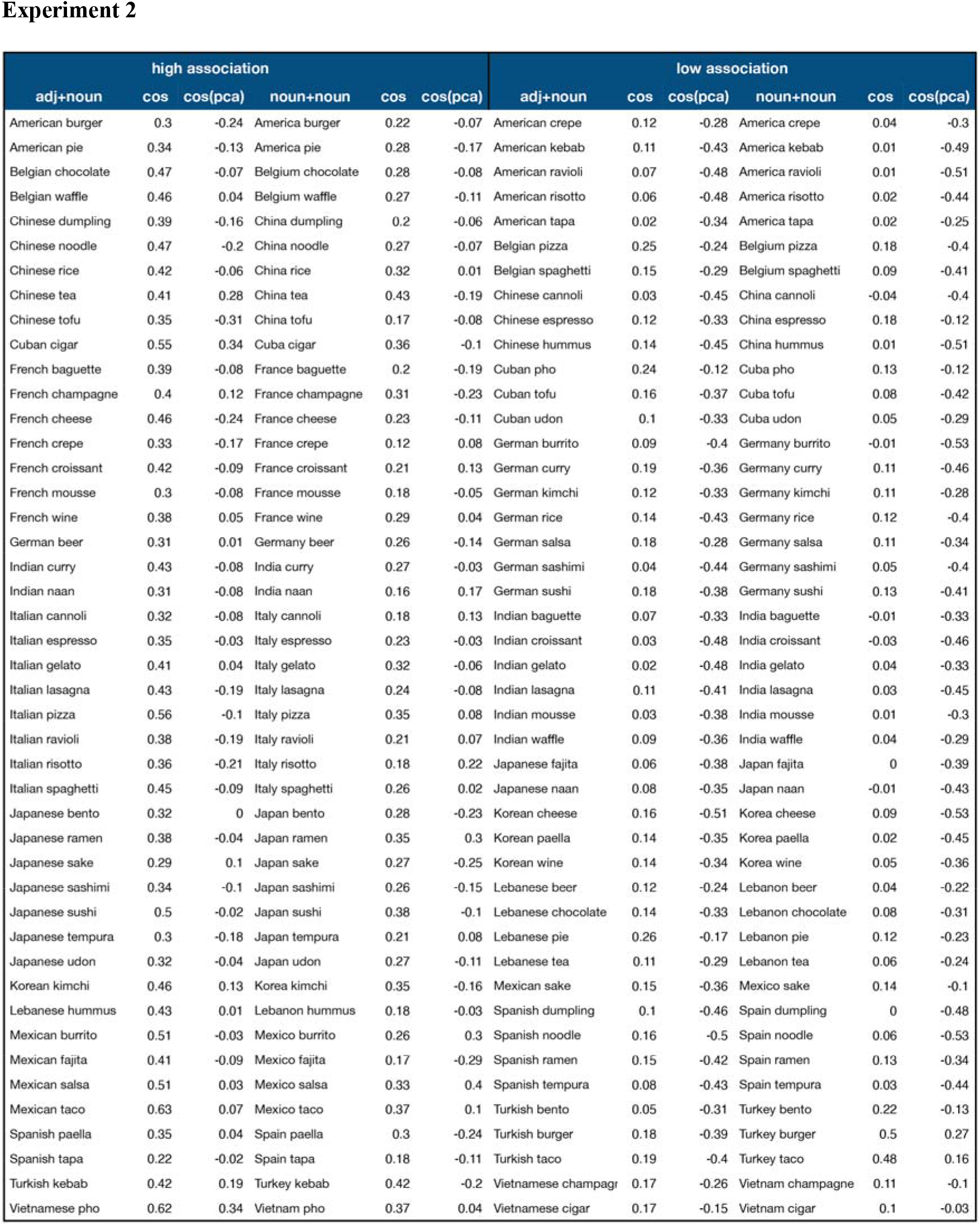
Stimuli lists for Experiment 1 and Experiment 2.

Experiment 1
| high association |  |  |  |  |  | low association |  |  |  |  |  |
| --- | --- | --- | --- | --- | --- | --- | --- | --- | --- | --- | --- |
| adj+noun | cos | cos(pca) | noun+noun | cos | cos(pca) | adj+noun | cos | cos(pca) | noun+noun | cos | cos(pca) |
| American burger | 0.39 | 0.15 | America burger | 0.2 | -0.07 | American crepes | 0.07 | -0.34 | America crepes | -0.01 | -0.37 |
| American pie | 0.39 | 0.02 | America pie | 0.2 | -0.17 | American dumplings | 0.09 | -0.52 | America dumplings | 0.07 | -0.46 |
| Belgian chocolate | 0.42 | 0.14 | Belgium chocolate | 0.21 | -0.08 | American kebab | 0.03 | -0.48 | America kebab | -0.03 | -0.46 |
| Belgian waffle | 0.43 | 0.11 | Belgium waffle | 0.27 | -0.11 | American paella | 0.17 | -0.34 | America paella | 0.11 | -0.42 |
| Chinese dumplings | 0.31 | 0.09 | China dumplings | 0.16 | -0.06 | American prawn | 0.08 | -0.39 | America prawn | -0.01 | -0.5 |
| Chinese noodles | 0.32 | 0 | China noodles | 0.18 | -0.07 | American risotto | 0.03 | -0.45 | America risotto | -0.04 | -0.41 |
| Chinese rice | 0.41 | 0.06 | China rice | 0.32 | 0.01 | American tapas | 0.02 | -0.47 | America tapas | 0.04 | -0.33 |
| Chinese tea | 0.43 | 0.01 | China tea | 0.24 | -0.19 | American tofu | 0.11 | -0.39 | America tofu | 0.03 | -0.43 |
| Chinese tofu | 0.52 | 0.15 | China tofu | 0.33 | -0.08 | American udon | 0.14 | -0.39 | America udon | 0.1 | -0.41 |
| Cuban cigar | 0.56 | 0.19 | Cuba cigar | 0.35 | -0.1 | Chinese cannoli | 0.19 | -0.33 | China cannoli | 0.14 | -0.33 |
| French baguette | 0.38 | 0.01 | France baguette | 0.21 | -0.19 | Chinese espresso | 0.06 | -0.4 | China espresso | 0.03 | -0.46 |
| French champagne | 0.36 | -0.05 | France champagne | 0.18 | -0.23 | English bibimbap | 0.06 | -0.49 | England bibimbap | 0.02 | -0.47 |
| French cheese | 0.45 | 0.11 | France cheese | 0.26 | -0.11 | English fajitas | 0.13 | -0.28 | England fajitas | 0.1 | -0.3 |
| French crepes | 0.32 | 0.1 | France crepes | 0.28 | 0.08 | English guacamole | 0.04 | -0.49 | England guacamole | 0.01 | -0.37 |
| French croissant | 0.29 | 0.08 | France croissant | 0.27 | 0.13 | English hummus | 0.1 | -0.35 | England hummus | 0.18 | -0.17 |
| French mousse | 0.34 | 0.03 | France mousse | 0.26 | -0.05 | English kimchi | 0.04 | -0.45 | England kimchi | 0.05 | -0.44 |
| French toast | 0.5 | 0.19 | France toast | 0.38 | 0.04 | English pho | 0.16 | -0.34 | England pho | 0.09 | -0.35 |
| French wine | 0.3 | -0.03 | France wine | 0.21 | -0.14 | English ravioli | 0.08 | -0.43 | England ravioli | 0.03 | -0.45 |
| German beer | 0.32 | 0 | Germany beer | 0.27 | -0.03 | English taco | 0.07 | -0.44 | England taco | 0.05 | -0.37 |
| Greek salad | 0.39 | 0.22 | Greece salad | 0.3 | 0.17 | English tequila | -0.01 | -0.35 | England tequila | 0 | -0.33 |
| Greek yogurt | 0.46 | 0.25 | Greece yogurt | 0.35 | 0.13 | English tortilla | 0.09 | -0.36 | England tortilla | 0.04 | -0.43 |
| Indian curry | 0.51 | 0.27 | India curry | 0.26 | -0.03 | German salsa | 0.06 | -0.38 | Germany salsa | 0.01 | -0.43 |
| Indian naan | 0.43 | 0.23 | India naan | 0.2 | -0.06 | German sashimi | -0.02 | -0.45 | Germany sashimi | 0 | -0.4 |
| Irish whiskey | 0.44 | 0.15 | Ireland whiskey | 0.21 | -0.08 | German yogurt | 0.04 | -0.42 | Germany yogurt | 0 | -0.44 |
| Italian cannoli | 0.51 | 0.31 | Italy cannoli | 0.33 | 0.08 | Greek cigar | 0.18 | -0.32 | Greece cigar | 0.11 | -0.42 |
| Italian cappuccino | 0.63 | 0.41 | Italy cappuccino | 0.37 | 0.07 | Indian baguette | 0.16 | -0.37 | India baguette | 0.13 | -0.42 |
| Italian espresso | 0.46 | 0.24 | Italy espresso | 0.37 | 0.22 | Indian cappuccino | 0.11 | -0.24 | India cappuccino | 0.08 | -0.31 |
| Italian gelato | 0.35 | 0.07 | Italy gelato | 0.3 | 0.02 | Indian croissant | 0.07 | -0.5 | India croissant | 0.07 | -0.39 |
| Italian lasagna | 0.31 | -0.06 | Italy lasagna | 0.18 | -0.23 | Indian gelato | 0.08 | -0.39 | India gelato | 0.07 | -0.42 |
| Italian latte | 0.62 | 0.53 | Italy latte | 0.37 | 0.3 | Indian lasagna | 0.11 | -0.27 | India lasagna | 0.09 | -0.3 |
| Italian pasta | 0.3 | -0.2 | Italy pasta | 0.22 | -0.25 | Indian mousse | 0.13 | -0.38 | India mousse | 0.09 | -0.41 |
| Italian pizza | 0.34 | -0.11 | Italy pizza | 0.28 | -0.15 | Indian vodka | 0.08 | -0.34 | India vodka | 0.15 | -0.41 |
| Italian ravioli | 0.47 | 0.28 | Italy ravioli | 0.28 | -0.1 | Indian waffle | 0.14 | -0.35 | India waffle | 0.08 | -0.29 |
| Italian risotto | 0.46 | 0.47 | Italy risotto | 0.27 | 0.08 | Irish bento | 0.09 | -0.32 | Ireland bento | 0.04 | -0.29 |
| Italian spaghetti | 0.41 | 0.06 | Italy spaghetti | 0.23 | -0.11 | Irish ramen | 0.12 | -0.48 | Ireland ramen | 0.05 | -0.51 |
| Japanese bento | 0.44 | 0.03 | Japan bento | 0.26 | -0.16 | Japanese naan | 0.14 | -0.45 | Japan naan | 0.08 | -0.48 |
| Japanese ramen | 0.42 | 0.05 | Japan ramen | 0.32 | -0.03 | Korean champagne | 0.15 | -0.21 | Korea champagne | 0.25 | -0.22 |
| Japanese sake | 0.41 | 0.15 | Japan sake | 0.43 | 0.3 | Korean cheese | 0.11 | -0.33 | Korea cheese | 0.06 | -0.26 |
| Japanese sashimi | 0.35 | -0.11 | Japan sashimi | 0.17 | -0.29 | Korean latte | 0.11 | -0.53 | Korea latte | 0.08 | -0.53 |
| Japanese sushi | 0.55 | 0.63 | Japan sushi | 0.36 | 0.4 | Korean whiskey | 0.1 | -0.28 | Korea whiskey | 0.08 | -0.15 |
| Japanese tempura | 0.4 | 0.12 | Japan tempura | 0.31 | 0.1 | Lebanese beer | 0.13 | -0.33 | Lebanon beer | 0.08 | -0.28 |
| Japanese udon | 0.46 | 0.08 | Japan udon | 0.23 | -0.24 | Lebanese chocolate | 0.16 | -0.5 | Lebanon chocolate | 0.09 | -0.56 |
| Korean bibimbap | 0.3 | -0.01 | Korea bibimbap | 0.18 | -0.11 | Lebanese tea | 0.03 | -0.39 | Lebanon tea | 0.01 | -0.3 |
| Korean kimchi | 0.38 | 0.07 | Korea kimchi | 0.18 | -0.2 | Russian burger | 0.07 | -0.23 | Russia burger | 0.12 | -0.36 |
| Lebanese hummus | 0.38 | 0.06 | Lebanon hummus | 0.29 | 0.04 | Russian burrito | 0.02 | -0.25 | Russia burrito | 0.14 | -0.25 |
| Mexican burrito | 0.31 | 0.07 | Mexico burrito | 0.26 | 0.01 | Russian curry | 0.12 | -0.28 | Russia curry | 0.04 | -0.25 |
| Mexican fajitas | 0.4 | -0.02 | Mexico fajitas | 0.14 | -0.39 | Russian laksa | 0.14 | -0.3 | Russia laksa | 0.13 | -0.56 |
| Mexican guacamole | 0.39 | 0.15 | Mexico guacamole | 0.16 | -0.12 | Russian pasta | 0.12 | -0.32 | Russia pasta | 0.07 | -0.39 |
| Mexican salsa | 0.47 | 0.34 | Mexico salsa | 0.28 | 0.09 | Russian spaghetti | 0.14 | -0.27 | Russia spaghetti | 0.11 | -0.22 |
| Mexican taco | 0.34 | -0.05 | Mexico taco | 0.24 | -0.08 | Scottish pizza | 0.05 | -0.39 | Scotland pizza | 0.06 | -0.27 |
| Mexican tequila | 0.35 | 0.03 | Mexico tequila | 0.23 | -0.03 | Scottish sushi | 0.12 | -0.33 | Scotland sushi | 0.18 | -0.08 |
| Russian vodka | 0.32 | -0.1 | Russia vodka | 0.22 | -0.14 | Singaporean pie | 0.13 | -0.36 | Singapore pie | 0.09 | -0.32 |
| Scottish haggis | 0.38 | 0.03 | Scotland haggis | 0.35 | 0 | Singaporean rice | 0.12 | -0.46 | Singapore rice | 0.08 | -0.41 |
| Singaporean laksa | 0.43 | 0.26 | Singapore laksa | 0.18 | -0.02 | Singaporean salad | 0.06 | -0.39 | Singapore salad | 0.01 | -0.47 |
| Spanish paella | 0.32 | 0.07 | Spain paella | 0.3 | 0.08 | Singaporean toast | 0.14 | -0.26 | Singapore toast | 0.12 | -0.21 |
| Spanish tapas | 0.48 | 0.46 | Spain tapas | 0.33 | 0.31 | Singaporean wine | 0.11 | -0.26 | Singapore wine | 0.07 | -0.24 |
| Spanish tortilla | 0.42 | 0.4 | Spain tortilla | 0.3 | 0.2 | Spanish haggis | 0.01 | -0.29 | Spain haggis | 0.01 | -0.29 |
| Thai prawn | 0.35 | 0.06 | Thailand prawn | 0.3 | 0.07 | Spanish noodles | 0.08 | -0.45 | Spain noodles | 0.06 | -0.4 |
| Turkish kebab | 0.4 | 0.12 | Turkey kebab | 0.21 | -0.16 | Spanish tempura | 0.06 | -0.47 | Spain tempura | 0 | -0.49 |
| Vietnamese pho | 0.42 | 0.24 | Vietnam pho | 0.42 | 0.21 | Turkish sake | 0.11 | -0.42 | Turkey sake | 0.01 | -0.5 |

**Table 1.** Stimuli lists for Experiment 1 and Experiment 2.
| high association |  |  |  |  |  | low association |  |  |  |  |  |
| --- | --- | --- | --- | --- | --- | --- | --- | --- | --- | --- | --- |
| adj+noun | cos | cos(pca) | noun+noun | cos | cos(pca) | adj+noun | cos | cos(pca) | noun+noun | cos | cos(pca) |
| American burger | 0.3 | -0.24 | America burger | 0.22 | -0.07 | American crepe | 0.12 | -0.28 | America crepe | 0.04 | -0.3 |
| American pie | 0.34 | -0.13 | America pie | 0.28 | -0.17 | American kebab | 0.11 | -0.43 | America kebab | 0.01 | -0.49 |
| Belgian chocolate | 0.47 | -0.07 | Belgium chocolate | 0.28 | -0.08 | American ravioli | 0.07 | -0.48 | America ravioli | 0.01 | -0.51 |
| Belgian waffle | 0.46 | 0.04 | Belgium waffle | 0.27 | -0.11 | American risotto | 0.06 | -0.48 | America risotto | 0.02 | -0.44 |
| Chinese dumpling | 0.39 | -0.16 | China dumpling | 0.2 | -0.06 | American tapa | 0.02 | -0.34 | America tapa | 0.02 | -0.25 |
| Chinese noodle | 0.47 | -0.2 | China noodle | 0.27 | -0.07 | Belgian pizza | 0.25 | -0.24 | Belgium pizza | 0.18 | -0.4 |
| Chinese rice | 0.42 | -0.06 | China rice | 0.32 | 0.01 | Belgian spaghetti | 0.15 | -0.29 | Belgium spaghetti | 0.09 | -0.41 |
| Chinese tea | 0.41 | 0.28 | China tea | 0.43 | -0.19 | Chinese cannoli | 0.03 | -0.45 | China cannoli | -0.04 | -0.4 |
| Chinese tofu | 0.35 | -0.31 | China tofu | 0.17 | -0.08 | Chinese espresso | 0.12 | -0.33 | China espresso | 0.18 | -0.12 |
| Cuban cigar | 0.55 | 0.34 | Cuba cigar | 0.36 | -0.1 | Chinese hummus | 0.14 | -0.45 | China hummus | 0.01 | -0.51 |
| French baguette | 0.39 | -0.08 | France baguette | 0.2 | -0.19 | Cuban pho | 0.24 | -0.12 | Cuba pho | 0.13 | -0.12 |
| French champagne | 0.4 | 0.12 | France champagne | 0.31 | -0.23 | Cuban tofu | 0.16 | -0.37 | Cuba tofu | 0.08 | -0.42 |
| French cheese | 0.46 | -0.24 | France cheese | 0.23 | -0.11 | Cuban udon | 0.1 | -0.33 | Cuba udon | 0.05 | -0.29 |
| French crepe | 0.33 | -0.17 | France crepe | 0.12 | 0.08 | German burrito | 0.09 | -0.4 | Germany burrito | -0.01 | -0.53 |
| French croissant | 0.42 | -0.09 | France croissant | 0.21 | 0.13 | German curry | 0.19 | -0.36 | Germany curry | 0.11 | -0.46 |
| French mousse | 0.3 | -0.08 | France mousse | 0.18 | -0.05 | German kimchi | 0.12 | -0.33 | Germany kimchi | 0.11 | -0.28 |
| French wine | 0.38 | 0.05 | France wine | 0.29 | 0.04 | German rice | 0.14 | -0.43 | Germany rice | 0.12 | -0.4 |
| German beer | 0.31 | 0.01 | Germany beer | 0.26 | -0.14 | German salsa | 0.18 | -0.28 | Germany salsa | 0.11 | -0.34 |
| Indian curry | 0.43 | -0.08 | India curry | 0.27 | -0.03 | German sashimi | 0.04 | -0.44 | Germany sashimi | 0.05 | -0.4 |
| Indian naan | 0.31 | -0.08 | India naan | 0.16 | 0.17 | German sushi | 0.18 | -0.38 | Germany sushi | 0.13 | -0.41 |
| Italian cannoli | 0.32 | -0.08 | Italy cannoli | 0.18 | 0.13 | Indian baguette | 0.07 | -0.33 | India baguette | -0.01 | -0.33 |
| Italian espresso | 0.35 | -0.03 | Italy espresso | 0.23 | -0.03 | Indian croissant | 0.03 | -0.48 | India croissant | -0.03 | -0.46 |
| Italian gelato | 0.41 | 0.04 | Italy gelato | 0.32 | -0.06 | Indian gelato | 0.02 | -0.48 | India gelato | 0.04 | -0.33 |
| Italian lasagna | 0.43 | -0.19 | Italy lasagna | 0.24 | -0.08 | Indian lasagna | 0.11 | -0.41 | India lasagna | 0.03 | -0.45 |
| Italian pizza | 0.56 | -0.1 | Italy pizza | 0.35 | 0.08 | Indian mousse | 0.03 | -0.38 | India mousse | 0.01 | -0.3 |
| Italian ravioli | 0.38 | -0.19 | Italy ravioli | 0.21 | 0.07 | Indian waffle | 0.09 | -0.36 | India waffle | 0.04 | -0.29 |
| Italian risotto | 0.36 | -0.21 | Italy risotto | 0.18 | 0.22 | Japanese fajita | 0.06 | -0.38 | Japan fajita | 0 | -0.39 |
| Italian spaghetti | 0.45 | -0.09 | Italy spaghetti | 0.26 | 0.02 | Japanese naan | 0.08 | -0.35 | Japan naan | -0.01 | -0.43 |
| Japanese bento | 0.32 | 0 | Japan bento | 0.28 | -0.23 | Korean cheese | 0.16 | -0.51 | Korea cheese | 0.09 | -0.53 |
| Japanese ramen | 0.38 | -0.04 | Japan ramen | 0.35 | 0.3 | Korean paella | 0.14 | -0.35 | Korea paella | 0.02 | -0.45 |
| Japanese sake | 0.29 | 0.1 | Japan sake | 0.27 | -0.25 | Korean wine | 0.14 | -0.34 | Korea wine | 0.05 | -0.36 |
| Japanese sashimi | 0.34 | -0.1 | Japan sashimi | 0.26 | -0.15 | Lebanese beer | 0.12 | -0.24 | Lebanon beer | 0.04 | -0.22 |
| Japanese sushi | 0.5 | -0.02 | Japan sushi | 0.38 | -0.1 | Lebanese chocolate | 0.14 | -0.33 | Lebanon chocolate | 0.08 | -0.31 |
| Japanese tempura | 0.3 | -0.18 | Japan tempura | 0.21 | 0.08 | Lebanese pie | 0.26 | -0.17 | Lebanon pie | 0.12 | -0.23 |
| Japanese udon | 0.32 | -0.04 | Japan udon | 0.27 | -0.11 | Lebanese tea | 0.11 | -0.29 | Lebanon tea | 0.06 | -0.24 |
| Korean kimchi | 0.46 | 0.13 | Korea kimchi | 0.35 | -0.16 | Mexican sake | 0.15 | -0.36 | Mexico sake | 0.14 | -0.1 |
| Lebanese hummus | 0.43 | 0.01 | Lebanon hummus | 0.18 | -0.03 | Spanish dumpling | 0.1 | -0.46 | Spain dumpling | 0 | -0.48 |
| Mexican burrito | 0.51 | -0.03 | Mexico burrito | 0.26 | 0.3 | Spanish noodle | 0.16 | -0.5 | Spain noodle | 0.06 | -0.53 |
| Mexican fajita | 0.41 | -0.09 | Mexico fajita | 0.17 | -0.29 | Spanish ramen | 0.15 | -0.42 | Spain ramen | 0.13 | -0.34 |
| Mexican salsa | 0.51 | 0.03 | Mexico salsa | 0.33 | 0.4 | Spanish tempura | 0.08 | -0.43 | Spain tempura | 0.03 | -0.44 |
| Mexican taco | 0.63 | 0.07 | Mexico taco | 0.37 | 0.1 | Turkish bento | 0.05 | -0.31 | Turkey bento | 0.22 | -0.13 |
| Spanish paella | 0.35 | 0.04 | Spain paella | 0.3 | -0.24 | Turkish burger | 0.18 | -0.39 | Turkey burger | 0.5 | 0.27 |
| Spanish tapa | 0.22 | -0.02 | Spain tapa | 0.18 | -0.11 | Turkish taco | 0.19 | -0.4 | Turkey taco | 0.48 | 0.16 |
| Turkish kebab | 0.42 | 0.19 | Turkey kebab | 0.42 | -0.2 | Vietnamese champagr | 0.17 | -0.26 | Vietnam champagne | 0.11 | -0.1 |
| Vietnamese pho | 0.62 | 0.34 | Vietnam pho | 0.37 | 0.04 | Vietnamese cigar | 0.17 | -0.15 | Vietnam cigar | 0.1 | -0.03 |

### Experiment procedures

Before recording, each subject’s head shape was digitized using a Polhemus dual source handheld FastSCAN laser scanner. Participants then completed the experiment while lying supine in a dimly lit, magnetically shielded room (MSR). At the Abu Dhabi facility, MEG data were recorded continuously using a whole-head 208 channel axial gradiometer system (Kanazawa Institute of Technology, Kanazawa, Japan); at the New York facility, MEG data were collected using a whole-head 156-channel axial gradiometer system (Kanazawa Institute of Technology, Kanazawa, Japan).

Stimuli were projected using PsychoPy2 (Peirce, 2019) onto a screen approximately 80 cm away from the participants’ eyes. The two words were presented for 300 ms each, in white 30-point Courier New font, on a gray background and subtended a vertical visual angle of 2°. Since high-associative phrases like “French wine” might be stored either as lexical items or as multi-word expressions (Arnon and Snider 2010; Jacobs et al., 2016; 2017), we presented the two words separately to encourage a combinatory or associative process instead of retrieving the phrase as a whole. An image appeared on screen after the target words and remained until subjects indicated whether it was a Match or a Mismatch to the preceding words, by pressing a button with the index finger of their left hand for a Match and middle finger for a Mismatch. No feedback was provided. This same picture-matching task has been employed in prior studies on semantic composition (e.g., Bemis and Pylkkänen, 2011; 2013). A blank screen was presented for 300 ms between each word and image. The inter-stimulus interval was normally distributed with a mean of 400 ms (SD=100 ms, min=135 ms, max=734 ms). Order of stimulus presentation was pseudo-randomized such that half of the trials were a Match and the other half a Mismatch. Each participant received a unique randomisation. Images for the categorization task were chosen to encourage concentration on both country nouns/adjectives and food nouns. For the two-word conditions, the target picture shows a food item with a country flag, and both the country and the food words were required to match; for the one-word condition, the picture contains only a food item. All images were color photographs found on Google Images. The whole recording session, including preparation time and practice, lasted around 40 minutes.

### MEG data acquisition and pre-processing

MEG data were recorded continuously at a sampling rate of 1000 Hz with an online 0.1 to 200 Hz band-pass filter. The raw data were first noise reduced via the Continuously Adjusted Least-Squares Method (Adachi et al., 2001) and low-pass filtered at 40 Hz. Independent component analysis (ICA) was then applied to remove artifacts such as eye blinks, heart beats, movements, and well-characterized external noise sources (mean ICA components=36; min ICA components=12; msc ICA components=66). The MEG data were then segmented into epochs spanning 100 ms pre-stimulus onset to 1200 ms post-stimulus onset. Epochs containing amplitudes greater than an absolute threshold of 2000 fT were automatically removed. Additional artefact rejection was performed through manual inspection of the data, removing trials that were contaminated with movement artefacts or extraneous noise. The whole epoch rejection procedure results in an average rejection rate of 9.4% (SD=6.5%) for each participant.

Cortically constrained minimum-norm estimates (mne; Hämäläinen and Ilmoniemi, 1994) were computed for each epoch for each participant. Following previous literature on semantic composition (e.g., Bemis & Pylkkänen., 2011;2013; Flick & Pylkkänen., 2020; Westerlund et al., 2015; Lyu et al., 2019), we used mne instead of whole-brain beamforming for source reconstruction. To perform source localization, the location of the participant’s head was coregistered with respect to the sensor array in the MEG helmet. For participants with anatomical MRI scans (n=5), this involved rotating and translating the digital scan to minimize the distance between the fiducial points of the MRI and the head scan. For participants without anatomical scans, the FreeSurfer (http://surfer.nmr.mgh.harvard.edu/) “fsaverage” brain was used, which involved first rotation and translation and then scaling the average brain to match the size of the head scan. A source space of 2562 source points per hemisphere was generated on the cortical surface for each participant. The Boundary Element Method (BEM) was employed to compute a forward solution, explaining the contribution of activity at each source to the magnetic flux at the sensors. Channel-noise covariance was estimated based on the 100 ms intervals prior to each artifact-free trial. The inverse solution was computed from the forward solution and the grand average activity across conditions. To lift the restriction on the orientation of the dipoles, the inverse solution was computed with “free” orientation, meaning that the inverse operator places three orthogonal dipoles at each location defined by the source space. When computing the source estimate, only activity from the dipoles perpendicular to the cortex were included. For each trial within one condition, the same inverse operator for that condition was applied to yield dynamic statistical parameter maps (dSPM) units (Dale et al., 1999) using an SNR value of 3. All data preprocessing steps were performed with MNE-python (v.0.19.2; Gramfort, et al., 2014) and Eelbrain (v.0.25.2; Brodbeck et al., 2014).

### Quantifying semantic associations

Semantic vectors for the country and food words were derived using the well-known GloVe word embeddings model (Pennington et al., 2014; freely available at https://nlp.stanford.edu/projects/glove/) trained on Common Crawl (https://commoncrawl.org/), which contains petabytes of raw web page data, metadata extracts and text extracts. GloVe model embodies the “distributional hypothesis” that words with similar meaning occur in similar contexts in an artificial neural network approach. Having obtained a 300-dimensional vector for each word, we then quantified how semantically associated each country word was to the food word by calculating the cosine similarity score between the two word vectors. Overall, our approach generated high associative phrases (e.g., “French/France cheese”) with a mean cosine similarity score of 0.34 (SD=0.1) and low associative pairs (e.g., “Korean/Korea cheese”) with a mean cosine similarity score of 0.08 (SD=0.05). Paired *t*-test revealed significant difference between the cosine similarity score of the two groups (*t*=29.8, *p*∼=0). To make the difference larger, we further applied principle component analysis (PCA) to the word embeddings and calculate the cosine similarity score for each word pair based on the first 30 PCs. This resulted in a mean cosine of 0.05 (SD=0.18) for the high-associative pairs and a mean cosine of -0.37 (SD=0.09) for the low-associative pairs. The group difference was significant (*t*=28.7, *p*∼=0), but it did not make the two groups more different (see Table 1 in Extended Data for the cosine similarity score for each word pair based on their 300 dimensional embeddings and the 30 PCs). Figure 1B visualizes all the word embeddings on a 2D scatter plot using t-SNE (van der Maaten, 2014).

## Statistical analysis

### Behavioral data analysis

Accuracy were analyzed using generalized linear mixed model (GLMM) with binomial error distribution and “logit” link, and response times (RTs) were analyzed using GLMM with lognormal transformation. For the maximal model, fixed effects included the main effects of composition, association and their interaction, as well as the differences between the single- and the average of all four two-word conditions. We included association as either a categorical or a continuous variable with the cosine similarity scores (both before and after PCA). Model comparison results showed that the optimal model including association as a categorical variable explained the behavioral data better than when association was included as a continuous variable. Random effects included by-subject random intercept and slopes of all fixed effects, as well as by-item random intercept (Barr et al., 2013). In addition, word one and word two frequency were used as control variables, which were calculated based on the combined word counts from the Google Book unigrams (http://books.google.com/ngrams) and the SUBTLEXus corpus (Brysbaert and New, 2009). The log frequency of each word is shown in Table 2 in Extended Data. RTs above or below than 3 standard deviations were removed under the assumption that these represented errors or distractions rather than task-related responses.

**Table 2.**
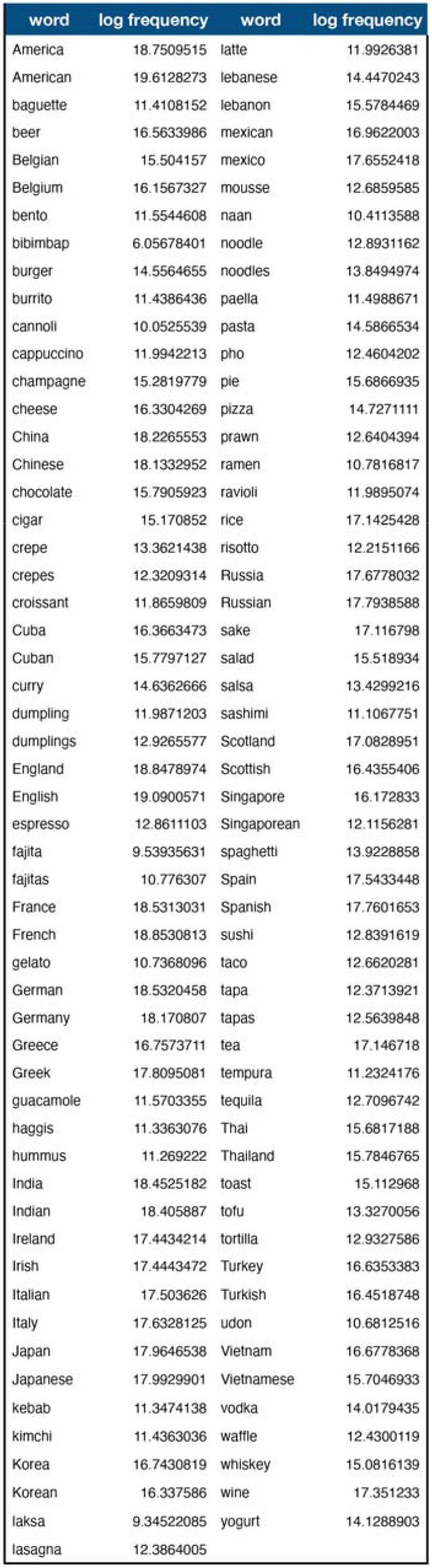
Log frequency for all the words in Experiment 1 and Experiment 2.

| word | log frequency | word | log frequency |
| --- | --- | --- | --- |
| America | 18.7509515 | latte | 11.9926381 |
| American | 19.6128273 | lebanese | 14.4470243 |
| baguette | 11.4108152 | lebanon | 15.5784469 |
| beer | 16.5633986 | mexican | 16.9622003 |
| Belgian | 15.504157 | mexico | 17.6552418 |
| Belgium | 16.1567327 | mousse | 12.6859585 |
| bento | 11.5544608 | naan | 10.4113588 |
| bibimbap | 6.05678401 | noodle | 12.8931162 |
| burger | 14.5564655 | noodles | 13.8494974 |
| burrito | 11.4386436 | paella | 11.4988671 |
| cannoli | 10.0525539 | pasta | 14.5866534 |
| cappuccino | 11.9942213 | pho | 12.4604202 |
| champagne | 15.2819779 | pie | 15.6866935 |
| cheese | 16.3304269 | pizza | 14.7271111 |
| China | 18.2265553 | prawn | 12.6404394 |
| Chinese | 18.1332952 | ramen | 10.7816817 |
| chocolate | 15.7905923 | ravioli | 11.9895074 |
| cigar | 15.170852 | rice | 17.1425428 |
| crepe | 13.3621438 | risotto | 12.2151166 |
| crepes | 12.3209314 | Russia | 17.6778032 |
| croissant | 11.8659809 | Russian | 17.7938588 |
| Cuba | 16.3663473 | sake | 17.116798 |
| Cuban | 15.7797127 | salad | 15.518934 |
| curry | 14.6362666 | salsa | 13.4299216 |
| dumpling | 11.9871203 | sashimi | 11.1067751 |
| dumplings | 12.9265577 | Scotland | 17.0828951 |
| England | 18.8478974 | Scottish | 16.4355406 |
| English | 19.0900571 | Singapore | 16.172833 |
| espresso | 12.8611103 | Singaporean | 12.1156281 |
| fajita | 9.53935631 | spaghetti | 13.9228858 |
| fajitas | 10.776307 | Spain | 17.5433448 |
| France | 18.5313031 | Spanish | 17.7601653 |
| French | 18.8530813 | sushi | 12.8391619 |
| gelato | 10.7368096 | taco | 12.6620281 |
| German | 18.5320458 | tapa | 12.3713921 |
| Germany | 18.170807 | tapas | 12.5639848 |
| Greece | 16.7573711 | tea | 17.146718 |
| Greek | 17.8095081 | tempura | 11.2324176 |
| guacamole | 11.5703355 | tequila | 12.7096742 |
| haggis | 11.3363076 | Thai | 15.6817188 |
| hummus | 11.269222 | Thailand | 15.7846765 |
| India | 18.4525182 | toast | 15.112968 |
| Indian | 18.405887 | tofu | 13.3270056 |
| Ireland | 17.4434214 | tortilla | 12.9327586 |
| Irish | 17.4443472 | Turkey | 16.6353383 |
| Italian | 17.503626 | Turkish | 16.4518748 |
| Italy | 17.6328125 | udon | 10.6812516 |
| Japan | 17.9646538 | Vietnam | 16.6778368 |
| Japanese | 17.9929901 | Vietnamese | 15.7046933 |
| kebab | 11.3474138 | vodka | 14.0179435 |
| kimchi | 11.4363036 | waffle | 12.4300119 |
| Korea | 16.7430819 | whiskey | 15.0816139 |
| Korean | 16.337586 | wine | 17.351233 |
| laksa | 9.34522085 | yogurt | 14.1288903 |
| lasagna | 12.3864005 |  |  |

If the maximal model could not converge, its random effects were orthogonalized, making the zero-correlation-parameter (ZCP) model. PCA was applied to the ZCP model results and random effects who explained less than 1% of total variances were removed from the ZCP model to make the reduced model. The extended model was built by adding back the correlations among random effects in the reduced model. If the extended model still could not converge, its random effects who explained less than 1% of total variances were further removed to make the updated extended model. This step was repeated until an extended model converge. Then the converged extended model was compared to the reduced model. The model, which explained the data better (with smaller Akaike’s Information Criterion) and used less parameters, was chosen as the optimal model whose results were reported (Bates et al., 2015).

The GLMM analyses were conducted via the “lme4” package (Bates et al, 2015) in R (v3.6.3) and RStudio (v1.2). Statistical significance of fixed effects was estimated using the “lmerTest” package (Kuznetsova et al., 2017), in which Satterthwaite’s approximation was applied to estimate degrees of freedom.

### Mass univariate multiple regression

We coded the composition and association factors as binary variables and performed a two-stage multiple regression with single-trial source estimates as dependent variables, and compositionality, strength of association and their interaction, and number of words as predictors (Figure 3A). We coded association as a binary variable as the behavioral analyses suggested that association as a categorical variable explains the response better than association as a continuous variable. We did not perform the same mixed-effect model for the behavioral analysis as the maximal model usually cannot converge, and we have to trim the random effects following Bates et al. (2015). Since we performed the mass univariate analysis for 755 sources*1301 timepoint, which means we have to conduct the trimming procedure for 755*1301= 982,255 times. This is not practical to implement. Analyzing global field power over time by source is not ideal either as we specifically want to know both the spatial and the temporal extent of our effects. Therefore we followed previous literature to use two-stage regression analyses for MEG data (e.g., Gwilliams et al., 2016)

Given the significant effects of word frequency on the behavioral results, we also included frequency of the first and second word as control variables. At the first stage, we applied an ordinary least squares regression for each subject’s single-trial source estimates for each source within a bilateral language mask at each millisecond from 100 millisecond before the onset of the first word to 1200 millisecond post-stimulus onset. The language mask (see the light pink region in Figure 4A and 4E) covered regions including the whole temporal lobe, the inferior frontal gyrus (IFG; defined as the combination of BAs 44 and 45), the ventromedial prefrontal cortex (vmPFC; defined as BA11), the angular gyrus (AG; defined as BA39) and the supramarginal gyrus (SMA; defined as BA 40). The left AG and vmPFC have also been implicated in previous literature on conceptual combination (Bemis and Pylkkänen, 2011; Price et al., 2015), and the LIFG and the LMTG have been suggested to underlie syntactic combination (Flick and Pylkkänen, 2020; Haggort, 2005; Lyu et al., 2019; Matchin et al., 2019; Matchin and Hicock, 2020). The first-stage regression resulted in a β coefficient for each variable at each source and each timepoint for each subject.

At the second stage, we performed a one-sample *t*-test on the distribution of the β values across subjects for each variable separately, again at each source and each timepoint, to test if their values were significantly different from zero. We applied the threshold-free cluster enhancement (TFCE) approach to identify significant spatiotemporal point within our mask (Smith and Nichols, 2009). The TFCE approach aims to enhance areas of signal that exhibit some spatial and temporal contiguity without relying on hard-threshold-based clustering. Each unthresholded *t*-statistic for a spatiotemporal point is passed through an algorithm which enhances the intensity within cluster-like regions. Precisely, the TFCE output for source *s* at time *t* is 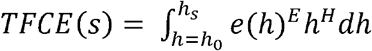, where *h*_*s*_ is the *t*-statistic value, *h*_*s*_ is 0, *e(h)* is the extent of the single cluster containing the point *p*. Therefore, each point’s TFCE score is the sum of the scores of all “supporting sections” underneath it, which is simply the height *h* (raised to some power *H*) multiplied by the cluster extent *e*(*h*) (raised to some power *E*). We repeated this procedure for 10,000 times. This involved randomly shuffling 0 and the β coefficient for each participant, repeating the mass univariate one-sample *t*-test and calculated the TFCE value for each spatiotemporal point within the mask and the analysis time window of 0-1200 ms after the stimulus onset. The observed TFCE value for each spatiotemporal point was subsequently assigned a *p*-value based on the proportion of random partitions that resulted in a larger test statistic than the observed one. Unlike the cluster-based approach, the TFCE approach allows us to make statistical inference about individual spatiotemporal point as each TFCE score has their own *p*-value.

### Multivariate pattern classification

To investigate more fine-grained encodings of the four conditions in the left anterior and middle temporal lobe, we conducted a searchlight multivariate pattern classification analysis within the same language mask used in the previous regression analysis. Under the assumption that patterns of brain activation contain information that distinguishes between the experimental conditions, we applied classification to 4 pairwise combinations of our 4 conditions: high-association phrases vs high-association lists, low-association phrases vs low-association lists, high-association phrases vs low-association phrases, and high-association lists vs low-association lists. The high-association phrases vs high-association lists classification and the low-association phrases vs low-association lists classification compares composition across the association levels, and the high-association phrases vs low-association phrases classification and the high-association lists vs low-association lists classification compares association across the composition levels.

We first regressed out the word frequency effects based on the regression analyses from the source estimate data, then we decimated the source estimates by a factor of 5, creating a 755 sources*260 timepoint matrix for each trial. We then combined the trials in each condition to 15 pseudo-trials by randomly dividing the trials into 15 partitions (4 trials per partition for Experiment 1 and 3 trials per partition for Experiment 2) and averaging across the trials within partitions. These two steps were performed to decrease the computational costs and increase the signal-to-noise ratio, as well as keeping the trial number the same across the two experiments (see Guggenmos et al., 2018).

We trained 4 linear Support Vector Machine (SVM) classifiers, each applied to a pairwise combination of the 4 conditions. The SVM is a widely used classification method due to its favorable characteristics of high accuracy, ability to deal with high-dimensional data and versatility in modeling diverse sources of data (Schölkopf et al., 2004). Linear SVM classifier was chosen as it has been shown to out-perform other classifiers on both MEG (Guggenmos et al., 2018) and fMRI data (Misaki et al., 2010). We performed a leave-one-stimulus-pair-out cross-validation procedure, where one randomly selected pseudo-trial was taken from each condition as a test sample and all remaining stimuli were used for classifier training. Classifier performance was estimated by averaging across the 100 permutations, and for each permutation we re-partitioned the trials to pseudotrials. The binary classifiers were separately applied to all spatiotemporal timepoints, with a radius of 100 sources * 5 timepoints.

The same analysis pipeline was applied to each subject. Classification accuracy averaged over subjects at each timepoint minus the chance level of 50% was submitted to a one-sample one-tailed *t*-test with TFCE correction for 10,000 permutations, where the sign of the *t-*statistic for each point was permuted. The analysis time window was between 600-1200 ms (Figure 3B). We used the python scikit-learn package (0.22.1) for the SVM analyses.

### Directed connectivity analysis

To further understand how LATL and LMTL exchange information during the two-word comprehension process, we conducted a directed connectivity analysis to estimate the temporal pattern of information flow between the two regions. The underlying logic is similar to that of the Granger causality analysis (Granger, 1969), namely, if the activity in region A has a causal effect on the activity in region B, then activity at a past time point of region A should contain information that helps predict current activity in region B above and beyond the previous activity in region B alone. Here we performed a multivariate pattern-based connectivity analysis which has been applied previously to estimate information flows between brain regions on both visual (Goddard et al., 2016) and language processing (Lyu et al., 2019).

We first calculated the data representational dissimilarity matrices (RDMs) of the two fROIs at time point *D(A, t)* and *D(B, t)* as 1 minus the pairwise Pearson *r*’s correlation among the conditions. The fROIs covered sources that are significant in either the univariate regression or the multivariate classification analysis. These RDMs indicate the degree to which different levels of associative strength and compositionality evoke similar or distinct response patterns in the neural population. To increase the signal-to-noise ratio, we adopted the same preprocessing procedures in the MVPA analysis, including regressing out word frequency effect, decimating the source estimates by a factor of 5 and combining the trials in each condition to 15 pseudo-trials. We then quantified the directed activity from region A to region B as the partial correlation coefficient between *D(A, t-dt)* and *D(B, t)*, partialling out *D(B, t−dt)*, where *dt* is the time interval between the current time point and the previous time point.

To avoid bias due to the choice of directionality and any specific previous time point, we computed the directed connectivity from region A to region B and from region B to region A for each time points from the onset of the second word to the following 600 ms, with a *dt* value of 5 ms before the current timepoint up till 600 ms (i.e., 5, 10, 15, …600 ms before the current time point). The extended time dimension allows us to determine the extent to which the current activity in the target region is correlated with the source region’s activity at each time point within the previous 600 ms. The same procedure was applied to each subject to obtain an average partial correlation coefficient matrix (Figure 3C). We report the results of this analysis in terms of the difference between the two directions of the information flow (A→B – B→A). Significance of the difference was determined by a one-sample *t*-test with 10,000 permutations.

### Data and code availability

Experimental stimuli and MEG single-trial source estimates for each participant is available to download via Open Science Framework (OSF) at https://osf.io/ea4s6/. All analyses were performed using custom codes written in Python, making heavy use of mne, eelbrain and scikit-learn. The analysis codes can be downloaded at https://osf.io/ea4s6/.

## Results

### Behavioral results

Overall, participants achieved a high accuracy of 90.9% (SD=28.2%) with a mean RT of 1.07 s (SD=0.77 s). The mixed-effects regression analysis revealed a significant effect of association for accuracy and a marginally significant effect for reaction time, where higher association between the two words increased accuracy (*t*=2.43, Cohen’s d=0.38, *p*=0.015) and reduced reaction time (*t*=-1.8, Cohen’s d=0.21, *p*=0.08). Composition was not significant for either accuracy (*t*=0.71, Cohen’s d=0.11, *p*=0.48) or reaction time (*t*=-0.72, Cohen’s d=0.11, *p*=0.48; see Figure 2A). We also observed a highly significant word frequency effect on both accuracy and RTs, such that more frequent first word reduced reaction time (*t*=-4.07, Cohen’s d=0.63, *p*∼=0), and more frequent second word both increased accuracy (*t*=6.19, Cohen’s d=0.95, *p*∼=0) and reduced reaction time (*t*=-8.3, Cohen’s d=1.28, *p*∼=0). This typical slowdown in responses for less frequent words indicates that the stimuli are being perceived as intended. Number of words is also significant for reaction time, such that single word conditions are faster than two word conditions (*t*=6.55, Cohen’s d=1.01, *p*∼=0; see Figure 2B).

**Figure 2.**
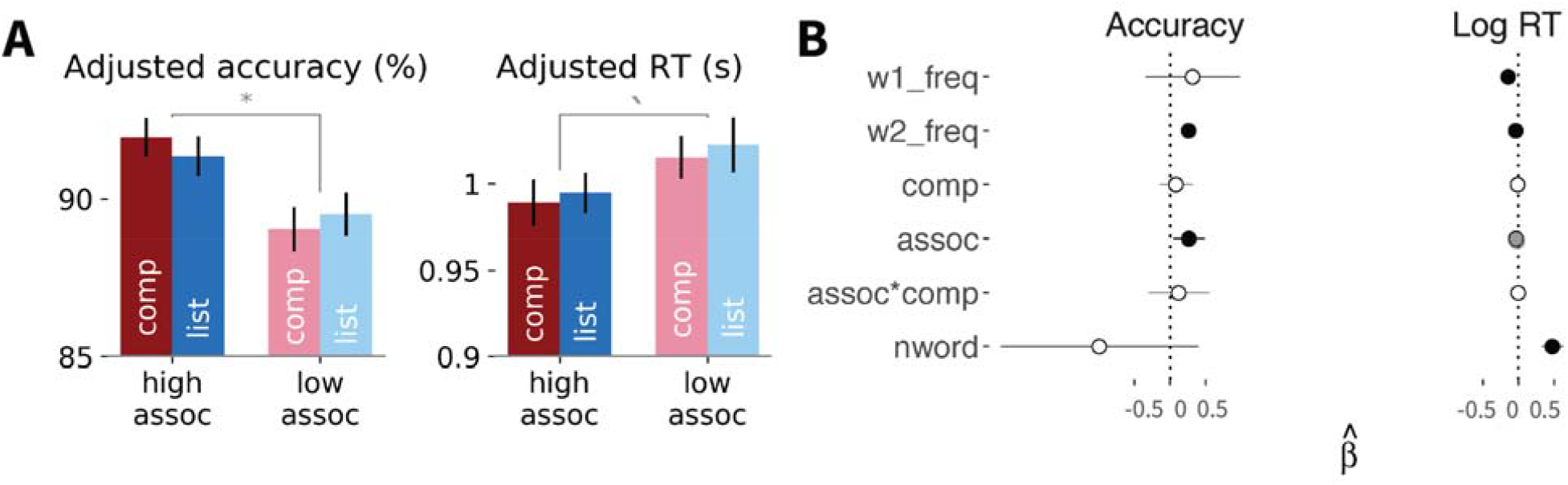
Behavioral results. ***A***, Mean predicted accuracy and reaction time for the four conditions, regressing out the effect of word frequency. Association is significant for accuracy (*p*=0.049) and marginally significant for reaction time (*p*=0.08). Error bars indicate 1 standard error. * *p*<.05. ***B***, Fitted coefficients for all predictors for accuracy and log reaction time. Error bars show 95% confidence intervals. Black point denotes significant coefficients at the level of *p*<0.05 and grey point denotes marginally significance at the level of *p*<0.1.

**Figure 3.**
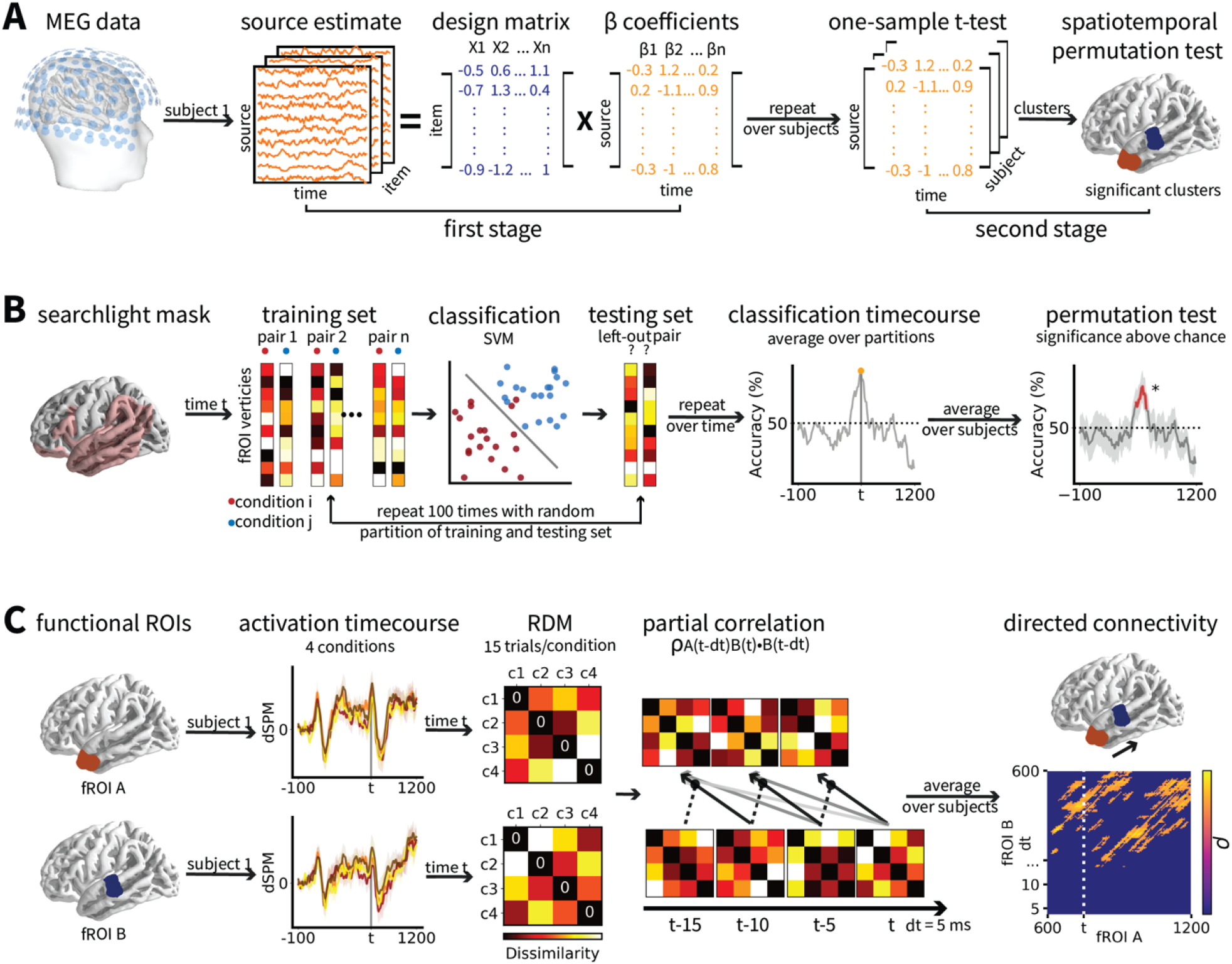
Schematic illustration of the analysis procedure. ***A***, Two-stage multiple regression analyses. At the first stage, an ordinary least squares regression was applied to each participant’s single-trial source estimates for each source within a selected region at each timepoint of the analysis window. At the second stage, a one-sample *t*-test was performed on the distribution of β values across subjects for each variable at each source and each timepoint, to test if their values were significantly different from zero. Significance were determined by TFCE correction with 10,000 permutations. ***B***, Searchlight multivariate pattern classification analyses. A linear Support Vector Machine (SVM) was trained on the combination of pseudo-trials from 2 conditions and tested on a left-out pair with 100 permutations. The same SVM analysis was applied independently to each source and timepoint within a language mask. Classification accuracy averaged over subjects at source and timepoint minus the chance level of 50% was submitted to a one-sample *t*-test and significance was determined by TFCE correction with 10,000 permutations. ***C***, Directed connectivity analyses. The representational dissimilarity matrices (RDMs) of the source estimates of the two fROIs derived from the regression and the classification analyses at time point *t* (*D(A, t)* and *D(B, t)*) were calculated as 1 minus the correlation between the conditions. Directed activity from region A to region B was quantified as the partial correlation coefficient between *D(A, t-dt)* and *D(B, t)*, partialling out *D(B, t−dt)*, where *dt* is the time interval between the current time point and the previous time point. Significance of coefficients greater than 0 was determined by 10,000 permutations with an alpha level of 0.05.

### Spatiotemporal clusters for association and composition

We identified one significant cluster for the interaction effect between association and composition from 637 to 759 ms (*t*=3.27, Cohen’s d=0.51, *p*=0.03), which is 37 ms after the onset of the second word till 159 ms afterwards. The cluster mainly covered the anterior temporal lobe (Figure 4A). The direction of the interaction is positive, which means that high-association compositional phrases elicited higher activity compared to other conditions (Figure 4B). We then calculated the mean fitted activation for the four conditions after controlling for the word frequency effect. The results showed that high-associative compositional phrases and low-associative lists were correlated with increased activity compared to low-associative compositional phrases and high-associative lists (Figure 4C,D).

**Figure 4.**
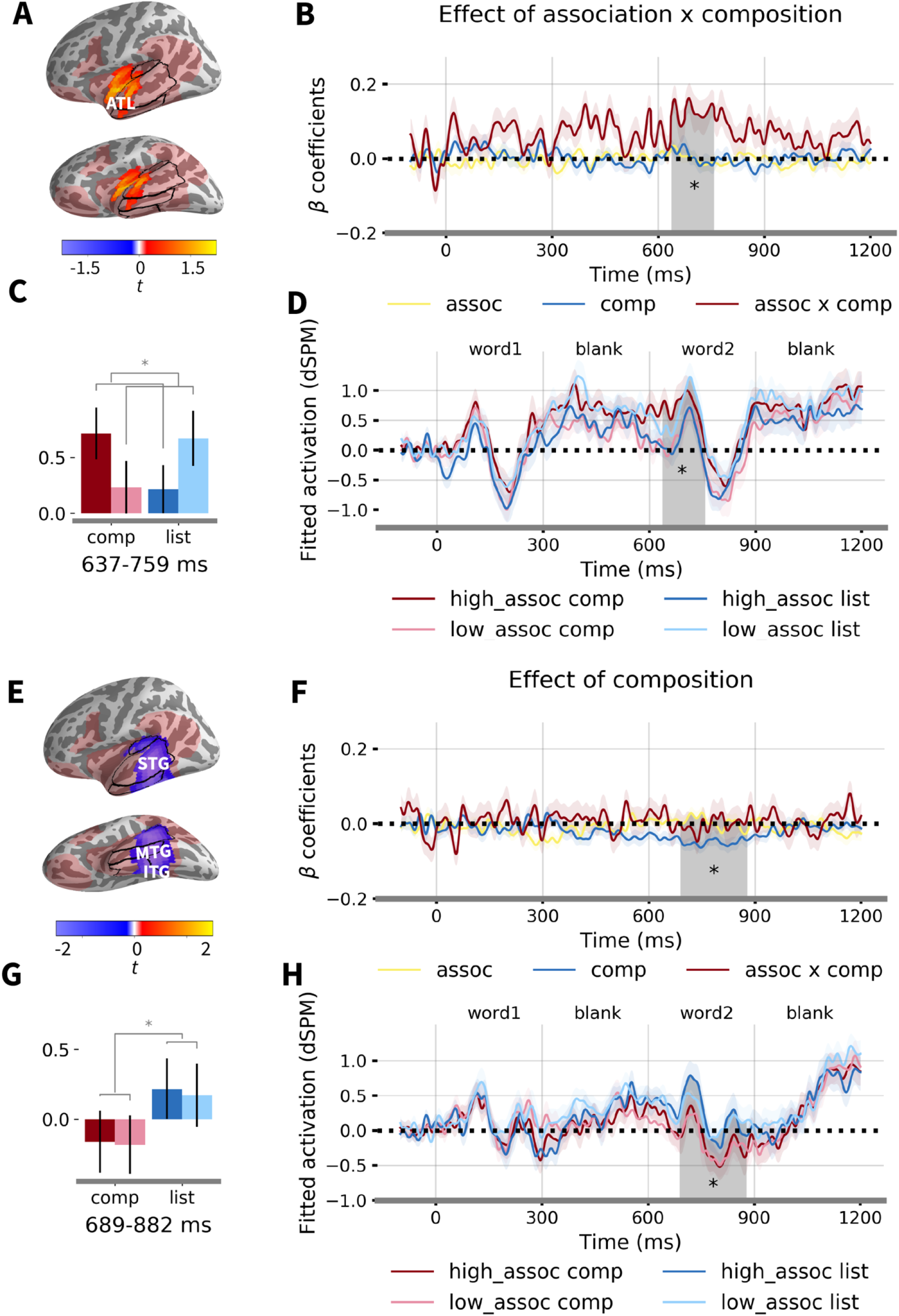
Multiple regression results. ***A***, Location of the significant cluster sensitive to the interaction effect between association and composition. Light red regions mark the language network within which the analysis was conducted. Color bar indicates *t*-statistics. ***B***, Timecourses of β coefficients averaged over the significant cluster. Shaded region denotes the significant time window from 637-759 ms (*p*=0.03). ***C***, Fitted responses for each condition averaged over the significant cluster and time window. ***D***, Timecourses of fitted response for each condition averaged over the significant cluster. Word frequency effects were regressed out of the responses. ***E***, Location of the significant cluster sensitive to the composition effect. ***F***, Timecourses of β coefficients averaged over the significant cluster. Shaded region indicates the significant time window from 689-882 ms (*p*=0.001). ***G***, Fitted responses for each condition averaged over the significant cluster and time window. ***H***, Timecourses of fitted response for each condition averaged over the cluster. Word frequency effects were regressed out of the responses. STG: superior temporal gyrus; MTG: middle temporal gyrus; ITG: inferior temporal gyrus; ATL: anterior temporal lobe.

The effect of composition was associated with one significant cluster in the middle temporal lobe from 689 to 882 ms (*t*=-4.05, Cohen’s d=0.63, *p*=0.001) after the onset of the first word (i.e., 89-282 ms after the onset of the second word). The cluster covered regions including the middle to posterior parts of the inferior, middle and superior temporal gyrus (Figure 4E). The sign of the β coefficient is negative, such that compositional phrase induced more activity with negative polarity (Figure 4F,G,H). No significant cluster was found for any of the effects in the right hemisphere within the mask, suggesting a highly left-lateralized neural activity for association and composition.

No significant cluster was found for the number of word predictor, suggesting that the two-word list and phrase conditions do not pattern together. We further conducted post-hoc pair-wise *t*-tests within the LATL and the LMTL clusters. We expected that the activation timecourses of the list conditions pattern with that of the single-word condition and this prediction was borne out: No significant temporal clusters were observed for the contrast between the two list conditions and the single word condition in either fROI. On the contrary, the two composition conditions were significantly different from the single-word condition in both the fROIs (see Figure 7).

Lexical frequency was significant in the middle temporal cortex from 202-294 ms (*t*=-4.11, Cohen’s d=0.64, *p*=0.034) for the first word and from 870-950 ms (i.e., 270-350 ms after the onset of the second word) for the second word (*t*=-4.72, Cohen’s d=0.74, *p*=0.046; see Figure 8). Here, higher word frequency decreased activation, consistent with most findings on the effect of frequency (Embick et al., 2001; Simon et al., 2017).

### Searchlight multivariate pattern classification

The classification results are shown in Figure 5. We observed on cluster in the LATL where classification accuracy for high-association composition vs. low-association composition was significantly higher than chance from 740-775 ms, that is, 140-175 ms after the onset of the second word (*t*=4.29, Cohen’s d=0.92, *p*=0.022). Accuracy of other classifiers was not significantly higher than chance. Compared with the multiple regression results, we observed the same distinction between high-association and low-association compositional phrases.

**Figure 5.**
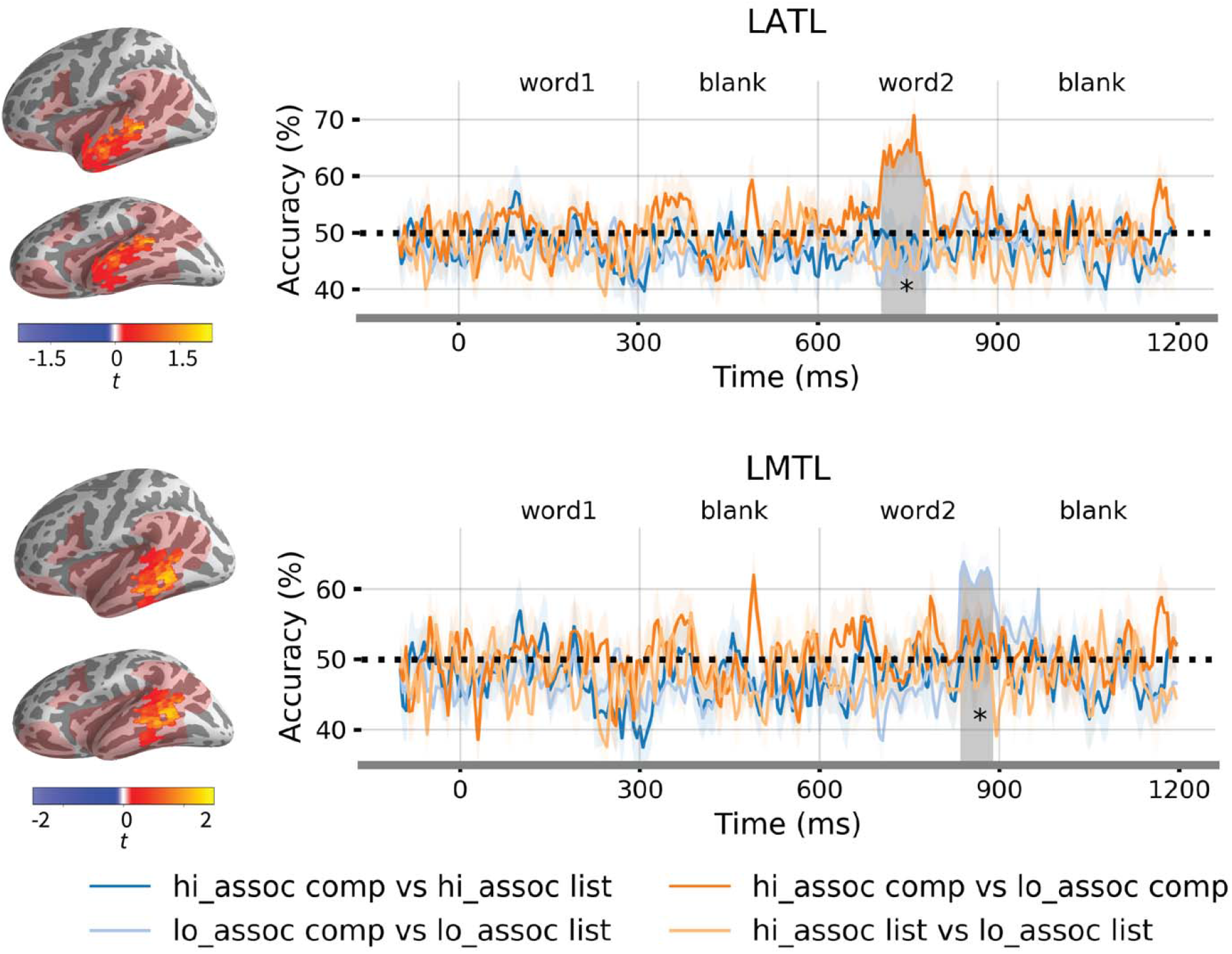
Searchlight multivariate pattern classification results. ***A***, Color-coded curves report classification timecourses for the 4 linear SVM classifiers averaged over the significant cluster in the LATL. Horizontal lines above the curves mark a statistically significant time window from 740-775 ms for the high-association composition vs low-association composition classifier (*p*=0.022). ***B***, Classification timecourses for the 4 linear SVM classifiers within the functional ROI for the composition effect identified by the regression analysis. Horizontal line above the curves marks a significant time window from 835-895 ms for the low-association composition vs low-association list classifier (*p*=0.016). Significance over the chance level of 50% was assessed by a permutation *t*-test with TFCE correction within the mask (light red region) and the analysis time window of 600-1200 ms.

The classifier for the low-association composition vs. low-association lists was significant in the LMTL at around 835-895 ms, i.e., 235-295 ms after the onset of the second word (*t*=5.55, Cohen’s d=0.87, *p*=0.016). Other classifiers did not perform significantly better than chance. This indicate that the effect of composition in the LMTL might be driven by the distinction between composition and list at the low-association level.

### Information flow between the active brain regions

Both out univariate regression analysis and the multivariate pattern classification have highlighted the LATL and the LMTL for high- and low-association compositional phrases, respectively. We further conducted a pattern-based directed connectivity analyses to disentangle the two effects at the temporal dimension. Comparison between the connectivity measures of the two directions revealed significant partial correlations from LATL to LMTL at around 0-250 ms after the onset of the second word. The significant time interval before the current timepoint ranged from the time intervals between 150 to 450 ms (*t*=6.26, Cohen’s d=0.98, *p*=0.03; see Figure 6A). Although the correlation value of ∼0.09 seems quite small, it is bigger than previous study using the same measures (Lyu et al., 2019). No significant partial correlations from LMTL to LATL was found by the contrast measure (Figure 6B). This suggests that information about associative strength and compositionality of the two words generated in the LATL within the previous 150-450 ms is continuously delivered to the adjacent LMTL from the onset of the second word to around 250 ms post-stimulus onset.

**Figure 6.**
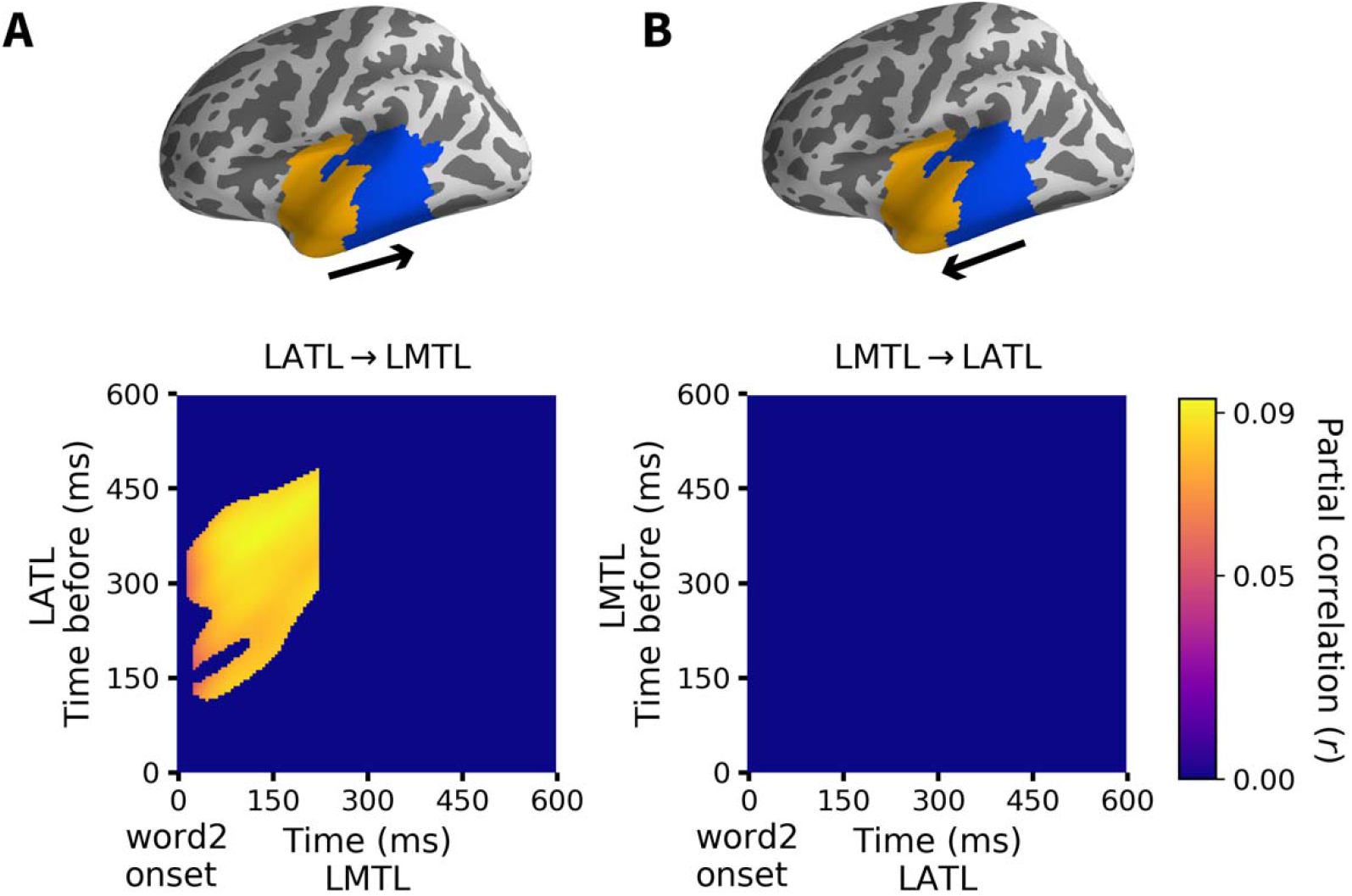
Directed connectivity between the LATL and the LMTL. ***A***, Contrast between the two directed connectivity measures show significant correlations from LATL to LMTL from around 0 to 250 ms after the onset of the second word, with a delay from about 150-450 ms (*p*=0.03). ***B***, No significant partial correlation coefficients matrix from LMTL to LATL.

Although the LATL and LMTL functional ROIs in our directed connectivity analysis are selected based on the regression results, this method is still largely data-driven as the connectivity strength is quantified by the partial correlations between the data RDMs. To investigate whether the connectivity results were specific to semantic association and composition or simply reflect intrinsic interactions between two adjacent regions, we conducted a supplementary control analysis with the primary somatosensory cortex (SMA, defined as BA1-3) and the primary motor cortex (M1; defined as BA4) in the left hemisphere as the two ROIs. Since the SMA and M1 were not related to either association or composition effect, the patterning of data RDMs in the two regions should bear no relation to our stimulus manipulation. Therefore, we expect no connectivity between the data RDMs of the two regions. The results supported our prediction: we did not find directed connectivity from SMA1 to M1 nor from M1 to SMA (see Figure 9).

**Figure 7.**
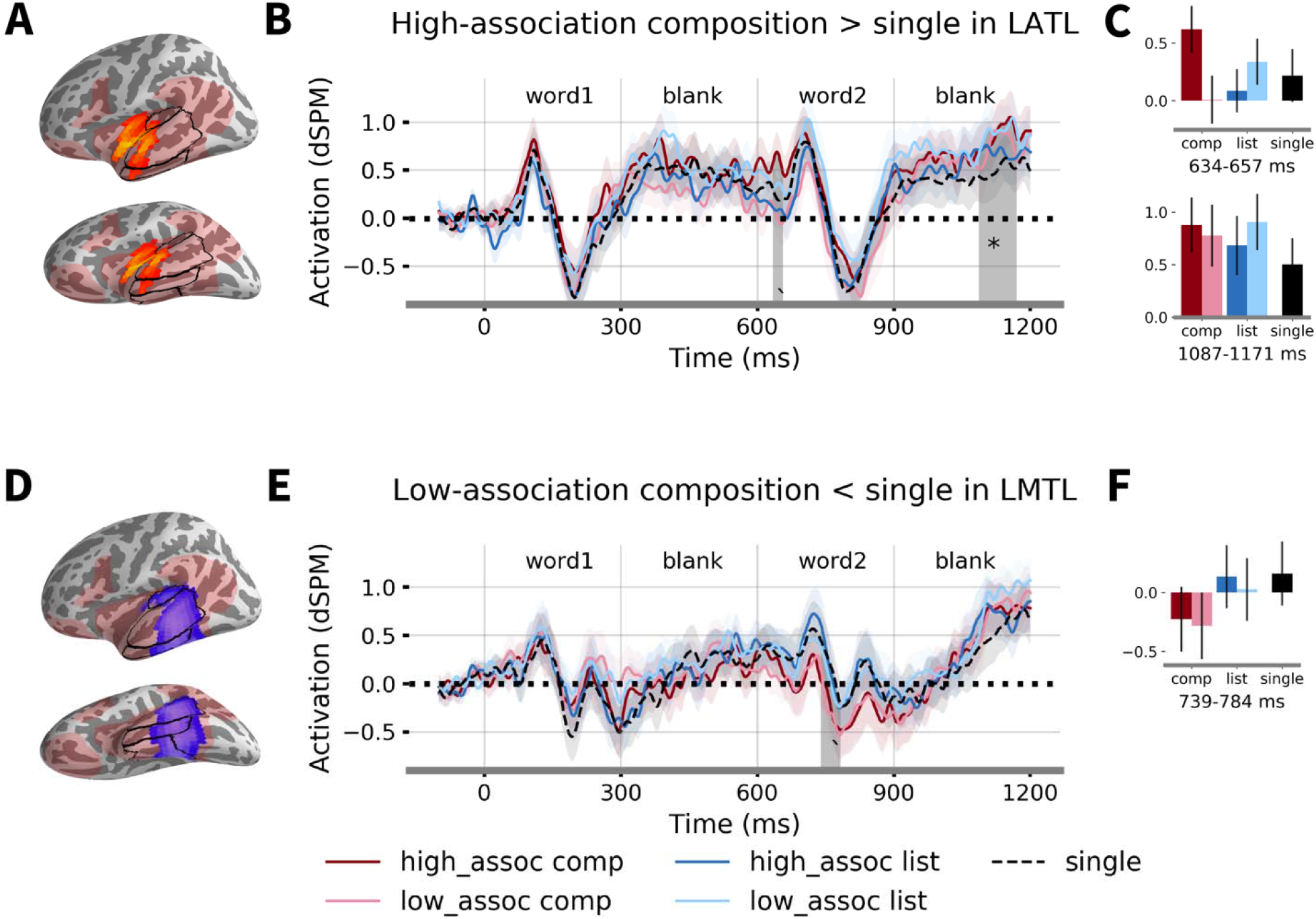
Pair-wise *t*-test results between each of the two-word conditions and the single word condition within the LATL and the LMTL fROIs. ***A***, The LATL fROI. ***B***, Time course of responses of for each condition averaged over the cluster. Shaded region indicates the marginally significant time windows of 634-657 ms (*p*=0.06) and significant time window of 1087-1171 ms for high-association composition > single (*p*=0.03. ***C***, Fitted responses for each condition averaged over the cluster and the time windows. ***D***, The LMTL fROI. ***E***, Time course of responses for each condition averaged over the cluster. Shaded region indicates the marginally significant time windows of 739-784 ms for low-association composition < single (*p*=0.08). ***F***, Fitted responses for each condition averaged over the cluster and the time window. Significance was determined by TFCE correction with 10,000 permutation. The testing time window is 600-1200 ms. * *p*<.05; ’ *p*<.1.

**Figure 8.**
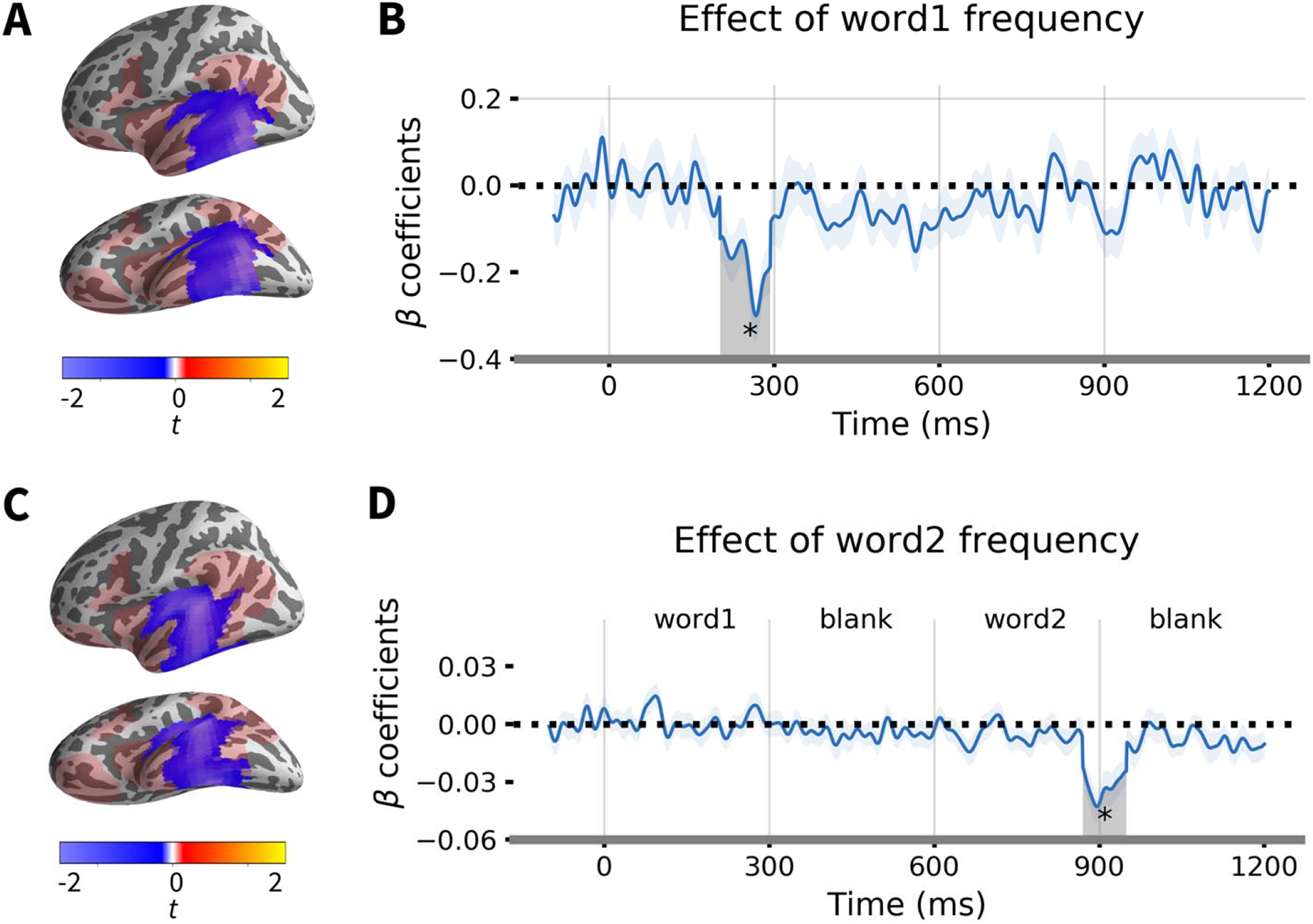
Effect of lexical frequency. ***A***, The significant cluster for the first word frequency. ***B***, Time course of beta coefficient of first word frequency averaged over the cluster. Shaded region denotes the significant time windows from 202-294 ms (*p*=0.034, TFCE corrected with 10,000 permutation). The analysis time window is 0-600 ms. ***C***, The significant cluster for the second word frequency. ***D***, Time course of beta coefficient of second word frequency averaged over the cluster. Shaded region denotes the significant time windows from 870-950 ms (*p*=0.045, TFCE corrected with 10,000 permutation). The analysis time window is 600-1200 ms.

**Figure 9.**
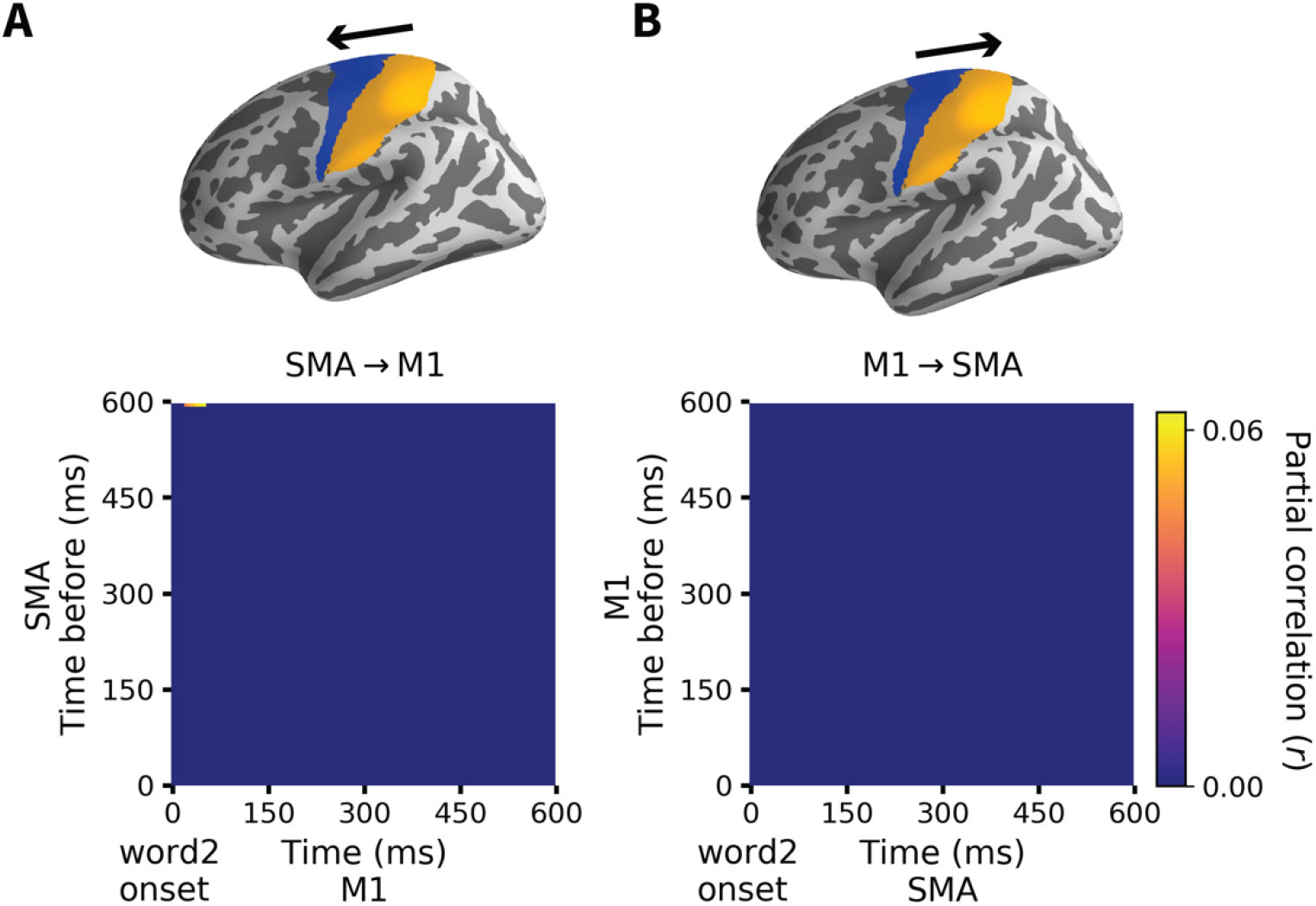
Directed connectivity between the left SMA and the left M1. ***A***, Partial correlation coefficients matrix from SMA to M1. ***B***, Partial correlation coefficients matrix from SMA to M1.

## Discussion

Prior work has identified similar brain regions for both semantic composition and associative encoding. Since combining meanings is intuitively different from simple association, this is somewhat surprising. Semantic compositionality allows human language to express an infinite number of complex expressions from their constituent meanings (Frege, 1980). This productive combinatory process is not dependent on the semantic relatedness of the constituent meanings. It engages even for totally novel combinations, such as “Cuban pho”. However, according to the distributional hypothesis of semantic theory, words that occur together are usually semantically related (see Boleda 2020 for a review). As famously put in Firth (1957): “a word is characterized by the company it keeps”, thus a common complex expression like “French cheese” involves both combinatory and associative processes. This raises the possibility that the two processes may have some type of common, neurally implemented core. We explored this possibility in the current work.

By orthogonalizing semantic composition and semantic association in our experimental design, we show that high-association compositional phrases elicited higher activity in the LATL than low-association compositional phrases. Searchlight multivariate pattern classification also shows distinctive activity patterns between the two conditions in the LATL. Thus there appears to be an early LATL process that reflects composition in a highly association-sensitive way. Note that it is possible that participants were thinking of adjective-noun phrases (e.g., “French cheese”) while seeing the noun-noun lists (e.g., “France cheese”) or construing the noun-noun lists as grammatical errors. These possibilities, however, could not explain the different activity patterns between the adjective-noun and noun-noun pairs at different association levels. If participants are more likely to think “French cheese” after seeing “France cheese” as compared to thinking “Korean cheese” after seeing “Korea cheese”, then we would argue that this is exactly because “France” and “cheese” are highly associated. Therefore, we think that our interaction effect in the LATL is best explained by the composition and association factors during the two-word comprehension.

Martin and Doumas (2019) suggested that meaning composition cannot be achieved in an association-only way such as the tensor product of constituent word embeddings, because human similarity judgements on “fuzzy cactus” and “fuzzy penguin” are not determined by the similarity between “cactus” and “penguin”. Martin & Doumas (2017) proposed a dynamic binding model (i.e., the symbolic-connectionist computational model DORA) that uses time to encode composition (see also Baggio, 2018 and Martin, 2020), yet it is unclear how such a model incorporates association between constituent words while interpreting compositional meanings. Given the interaction effect of association and composition in the LATL, we suggest that a cognitive model for meaning composition should include both association and composition factors.

The LATL’s sensitivity to both composition and association supports the role of both compositional and distributional semantics, which has been the subject of much discussion in Natural Language Processing and cognitive science. Distributional semantic models work by building co-occurrence vectors for every word in a corpus based on its context. They provide concrete information of word meaning but do not scale up to larger constituents of text, such as phrases or sentences. Compositional formal semantics complements this approach and various models have been proposed to combine the two paradigms (e.g., Mitchell and Lapata, 2008). These models typically use one algebraic compositional function such as addition or multiplication over constituent word embeddings to achieve composition. Alternatively, recent deep neural networks have been shown to capture grammar-dependent composition without clear-cut, algebraic rules (see Baroni, 2019). Here we show that the LATL responded differently to high- and low-association compositional phrases, suggesting that the neural mechanism of composition may indeed not rely on systematic compositional rules.

At the temporal dimension, the regression analysis showed an early LATL activity for the interaction effect of association and composition after the onset of the second word. This effect even shows up before the onset of the second word (544-846 ms) under a cluster-based permutation test (Maris and Oostenveld, 2007). However, in the cluster-based permutation test, the spatiotemporal cluster is formed before the permutation test, thus we cannot claim statistical significance for individual spatiotemporal points in the cluster. We therefore used the TFCE approach which allows us to infer the significance of each spatial and temporal point. The TFCE results showed a narrower spatiotemporal cluster in extent, where the significant time window is from 637-759 ms (i.e., 37-159 ms after the onset of the second word). This early onset likely reflects the high predictability of the high-associative stimuli, that is, some of the noun-meanings may be preactivated during the processing of the first word. For instance, when seeing “France/French”, participants might automatically think of French-related food, even though the second word could be “kimchi”.

In the LMTL, the regression analysis and the multivariate pattern classification results indicated that this region is mainly sensitive to the composition at the low-association level, i.e., the distinction between “Korean cheese” and “Korea cheese”. Compared to high-association compositional phrases, low-association compositional phrases form a newly-encountered meaning, hence likely involve more syntactic processing for a successful parsing. There is ample evidence that the LMTL supports syntactic combination (e.g., Flick and Pylkkänen, 2020; Lyu et al., 2019; Matchin et al., 2019; Matchin and Hicock, 2020). For example, the left pMTG in verb-noun combination shows the strongest model fit in the left middle temporal gyrus (Lyu et al., 2019), and both noun and verb phrases (e.g., “the frightened boy” and “frightened the boy”) induced greater activity in the superior temporal sulcus compared to word lists (Matchin et al., 2019). Moreover, post-nominal adjective-noun composition elicited activity in the left medial to posterior temporal lobe when semantics is held constant (Flick and Pylkkänen, 2020). Our composition effect in the LMTL might be attributed to the effect of word type (adjective vs noun; see Mollo et al., 2018 for distinction of noun and verb context in the LMTL). However, word type cannot explain the difference between the high-associative and low-associative compositional phrases in the LATL, since both “French cheese” and “Korean cheese” are grammatical. Note that we are not suggesting that the LMTL is only involved in syntactic processing. As shown in Figure 8, lexical frequency of both the first and second word was significantly correlated with the LMTL activity, such that higher frequency led to decreased activation at around 200-350 ms after the word onset. This spatiotemporal cluster is consistent with previous findings on the effect of frequency (e.g., Embick et al., 2001; Simon et al., 2017). By adding the first and second word frequencies as control variables in our regression model, we revealed the effect of composition in the LMTL in addition to the word frequency effect.

Our findings align with the controlled semantic cognition (CSC) framework (Jefferies, 2013; Lambon Ralph et al., 2017), which proposes a functional dissociation of the anterior and middle temporal regions as well. Under this framework, the ATL integrates information from different sources into a coherent concept, whereas the posterior middle temporal gyrus (pMTG) retrieves and manipulates semantic knowledge, such as retrieving infrequent semantic associations. Previous studies using a similar two-word MEG paradigm, showed that the ATL responded strongly to semantically coherent word pairs, whereas the posterior MTG was more sensitive to weak associations (Teige et al., 2018; 2019). Additionally, studies on semantic disorders also showed different patterns of impairment: patients with focal ATL atrophy exhibit symptoms of semantic dementia (SD), with performance strongly influenced by concept familiarity (Ding et al., 2020). In contrast, patients with temporoparietal damage are usually diagnosed with semantic aphasia (SA), characterized by their deficits in controlled semantic retrieval (Hoffman et al., 2018; Jefferies and Lambon Ralph., 2006; Jefferies et al., 2020). Mirroring Teige et al.’s (2018, 2019) findings and the SD/SA patients’ profiles, we also observed the LATL activation for semantically related word pairs, and the LMTL response to infrequent semantic associations. However, we added one important element to the converging story about the role of the LATL in semantic processing: syntactic knowledge. Unlike Teige et al.’s experiments which used only noun-noun word lists as stimuli, we also included meaning-controlled adjective-noun phrases. We showed that both composition and association contributed to semantic processing in the LATL, in line with a different stream of literature that has shown evidence for the LATL’s role in conceptual composition (e.g., Bemis and Pylkkänen, 2011; Pylkkänen, 2019). Compared to Teige et al.’s (2018, 2019) findings that association effect starts from around 200-400 ms after the onset of the target word, our interaction effect between association and composition began within 100 ms after the onset of the second word. This earlier interaction effect suggests that syntactic knowledge further facilitates efficient retrieval of highly coherent semantic information in the ATL under the CSC framework.

To directly probe the information flow between the two regions, we further conducted pattern-based directed analysis and showed that activity originated in the LATL 150-450 ms before it was continuously delivered to the LMTL within 250 ms after the stimulus onset. Prior studies have also suggested functional connectivity between the LATL and the LMTL in semantic association judgements (e.g., Jackson et al., 2016), and the LATL and the LMTL are also part of the default mode network, which has been shown to respond strongly when the probe and target items were highly overlapping conceptually (Lanzoni et al., 2020; Wang et al., 2020). Our results provide novel evidence on the direction of the functional connection between the LATL and the LMTL. Since our regression and decoding analyses showed that the LATL is more sensitive to the high-association compositional phrases while the LMTL is responding to the low-association compositional phrases, we therefore infer that the information flow from the LATL to the LMTL suggests that high-association composition precedes low-association composition. We interpreted these results as reflecting a functional dissociation where the LATL supports “shallow” meaning composition of common phrases, while the LMTL is engaged to interpret novel phrases via syntactic composition. At the temporal dimension, “shallow” meaning formation occurs earlier in the LATL, and triggers syntactic composition in the LMTL when the complex meaning is unfamiliar. Note that the anterior temporal region might be a “higher” cognitive region than the posterior temporal region since under the dual-stream model of language processing (Hickok and Poeppel, 2007), the posterior temporal regions precede the anterior temporal regions in the ventral pathway, and the posterior temporal regions correspond to the lexical interface which links phonological and semantic information, whereas the more anterior locations correspond to the “higher level” combinatorial network. In addition, the ATL is also considered a “hub” that integrates information from modality-specific regions under the “hub-and-spoke” model of semantic knowledge (Patterson et al., 2007). Thus the direction of flow from the LATL to the LMTL in our directed connectivity analysis likely reflects the flow-back of information from the higher to the lower cognitive regions.

In summary, we show evidence for distinct profiles in the left temporal lobe where the LATL is mostly sensitive to high-association compositional phrases, while the LMTL responds more to low-association compositional phrases. We also observed a prominent information flow from the LATL to the LMTL, suggesting that the integration of adjective and noun properties originated earlier in the LATL is consistently delivered to the LMTL when the complex meaning is newly encountered.

## Acknowledgments

This work was supported by the NYU Abu Dhabi Institute under Grant G1001. We thank Haiyan Jin for help with the statistical analysis on the behavioral data. We thank the reviewers for their invaluable comments and suggestions on earlier drafts of the paper.

